# Reductive evolution and unique infection and feeding mode in the CPR predatory bacterium *Vampirococcus lugosii*

**DOI:** 10.1101/2020.11.10.374967

**Authors:** David Moreira, Yvan Zivanovic, Ana I. López-Archilla, Miguel Iniesto, Purificación López-García

## Abstract

The Candidate Phyla Radiation (CPR) constitutes a large supergroup of mostly uncultured bacterial lineages discovered through metabarcoding and metagenomics in diverse environments. Having small cell sizes, reduced genomes, and limited biosynthetic capabilities, they are thought to be symbionts of other organisms from which they obtain essential biomolecules. However, the nature of this symbiosis (mutualistic, neutral, or parasitic) has been ascertained only for rare cultured members of the CPR phylum Saccharibacteria, which are epibiotic parasites of other bacteria. Here, we characterize the biology and the genome of *Vampirococcus lugosii*, which becomes the first described species of *Vampirococcus*, a genus of epibiotic bacteria morphologically identified decades ago. *Vampirococcus* belongs to the CPR phylum Absconditabacteria. It feeds on anoxygenic photosynthetic gammaproteobacteria, fully absorbing their cytoplasmic content. The cells divide epibiotically, forming multicellular stalks whose apical cells can more easily reach new hosts. *Vampirococcus* genome is small (1.3 Mbp) and highly reduced in biosynthetic metabolism genes. However, it is enriched in genes related to an elaborate, fibrous cell surface likely involved in complex interactions with the host. Comparative genomic analyses show that gene loss has been continuous during Absconditabacteria, and generally most CPR bacteria, evolution. Nonetheless, gene loss was compensated by gene acquisition by horizontal gene transfer and evolution *de novo*. In *Vampirococcus*, these innovations include new CRISPR-Cas effectors and a novel electron transport chain. Our findings confirm parasitism as a widespread lifestyle of CPR bacteria, which probably play a previously neglected virus-like ecological role in ecosystems, controlling bacterial populations by a unique form of predation.

## Introduction

Most prokaryotic organisms have remained unknown to science for a long time because of their limited morphological diversity and our inability to culture them. However, over the past few decades, the increasing use of molecular methods opened the black box of prokaryotic diversity^1^. Metabarcoding based on 16S rRNA gene PCR-amplification and sequencing (via, initially, Sanger and, more recently, diverse high-throughput sequencing techniques), unveiled a huge diversity of prokaryotes across ecosystems, including many high-rank (e.g. phylum or class level) clades for which we still lack cultured representatives. Direct, PCR-independent, massive sequencing of environmental DNA (metagenomics) completed this picture by i) revealing lineages whose rRNA genes escaped PCR amplification and ii) providing data about the complete gene complement of microbial communities and, hence, about their metabolic potential^1–4^. The possibility to assemble genome sequences from single cell amplified genomes^5^ or by binning from complex metagenomes^6^ has further led to gain genome-based knowledge for specific uncultured groups. Some of these groups are widely diverse and/or have pivotal importance in evolution, such as the eukaryote-related Asgard archaea^7^. Another such group is the ‘Candidate Phyla Radiation’ (CPR), which encompasses several dozens of high-rank lineages that represent a substantial diversity of the bacterial domain^2,8^.

Genome-resolved metagenomics has allowed the reconstruction of complete genome sequences for many CPR lineages. Despite the impressive diversity of this group, a clear common pattern distills from these sequences: all CPR bacteria contain small genomes (often <1 Mb) that encode limited metabolic capacities^3,8,9^. Most CPR bacteria are inferred to be obligatory anaerobes that ferment different substrates, lacking aerobic or anaerobic respiration. In most cases, their small genomes also lack genes involved in basic biosynthetic capacities, notably of lipids, nucleotides, and most amino acids^3,8^. These characteristics mirror those found in the DPANN archaea, which also exhibit small genomes and simplified metabolisms^9,10^. Consistent with their reduced genomes, CPR bacteria consist of very small cells when observed under the microscope, and are often able to pass through 0.2 μm pore-diameter filters^11^. The cultivation of the first species belonging to the CPR clade (specifically to the phylum Saccharibacteria, previously known as TM7) showed its cells growing attached to the surface of the human-associated actinobacterium *Actinomyces odontolyticus*^12^. More recently, three additional Saccharibacteria species growing in epibiotic association with diverse Actinobacteria of the human oral microbiome have been isolated^13^. However, although there is increasing support for the ultra-small CPR bacteria being epibionts that depend on hosts with more complete biosynthetic repertoires, current evidence remains fragmentary and based on these very few documented species of Saccharibacteria. Important questions such as the generality and nature (beneficial, neutral or deleterious) of CPR bacteria-host interactions need to be addressed on additional species to get global insight on the biology of this bacterial supergroup.

Here, we present the in-depth characterization of *Candidatus* Vampirococcus lugosii, a new CPR species from the phylum Absconditabacteria (previously known as SR1) found in the athalassic salt lake Salada de Chiprana. The genus *Vampirococcus* was described several decades ago as one of the rare examples of predatory bacteria^14^, but its phylogenetic identity remained elusive until now. Our observation of living plankton from this lake showed that *Ca*. V. lugosii grows and characteristically divides multiple times attached to cells of its specific host, the anoxygenic photosynthetic gammaproteobacterium *Halochromatium* sp., until it completely absorbs its cytoplasmic content thereby killing it. *Ca*. V. lugosii has a small genome (1.3 Mb), coding for an extremely limited biosynthetic metabolism but for an elaborate cell surface, most likely involved in a complex interaction with the host. These findings suggest that parasitism is widespread in CPR bacteria and that they play a previously neglected virus-like ecological role in controlling bacterial populations in many ecosystems.

## Results and Discussion

### Identification of ultra-small cells associated with blooms of anoxygenic phototrophic gammaproteobacteria

The Salada de Chiprana (NE Spain) is the only permanent athalassic hypersaline lake in Western Europe. It harbors thick, conspicuous microbial mats covering its bottom (Fig. 1a-b) and exhibits periodic stratification during which the deepest part of the water column becomes anoxic and sulfide-rich, favoring the massive development of sulfide-dependent anoxygenic photosynthetic bacteria^15^. We collected microbial mat fragments that were maintained in culture in the laboratory. After several weeks, we observed a bloom of anoxygenic photosynthetic bacteria containing numerous intracellular sulfur granules (Fig. 1c). Many of these bacterial cells showed one or several much smaller and darker cells attached to their surface (Fig. 1d-e). These cells were highly mobile, swimming at high speed with frequent changes of direction (Supplementary Video 1), in contrast with the non-infected cells that displayed slower (approximately half speed) and more straight swimming. Sometimes, two or more photosynthetic cells were connected through relatively short filaments formed by stacked epibiont cells (Fig. 1f). Although the photosynthetic cells carrying these epibionts were often alive and actively swimming, in some cases epibionts were associated to empty ghost cells where only the photosynthesis-derived sulfur granules persisted. Closer scrutiny of the epibionts revealed that they actually consisted of piles of up to 10 very flattened cells of 500-600 nm diameter and 200-250 nm height (e.g., Fig. 1d). These characteristics (size, morphology, and specific attachment to sulfide-dependent anoxygenic photosynthetic bacteria) perfectly matched the morphological description of the genus *Vampirococcus* observed over forty years ago in sulfidic freshwater lakes^14^.

**Fig. 1.**
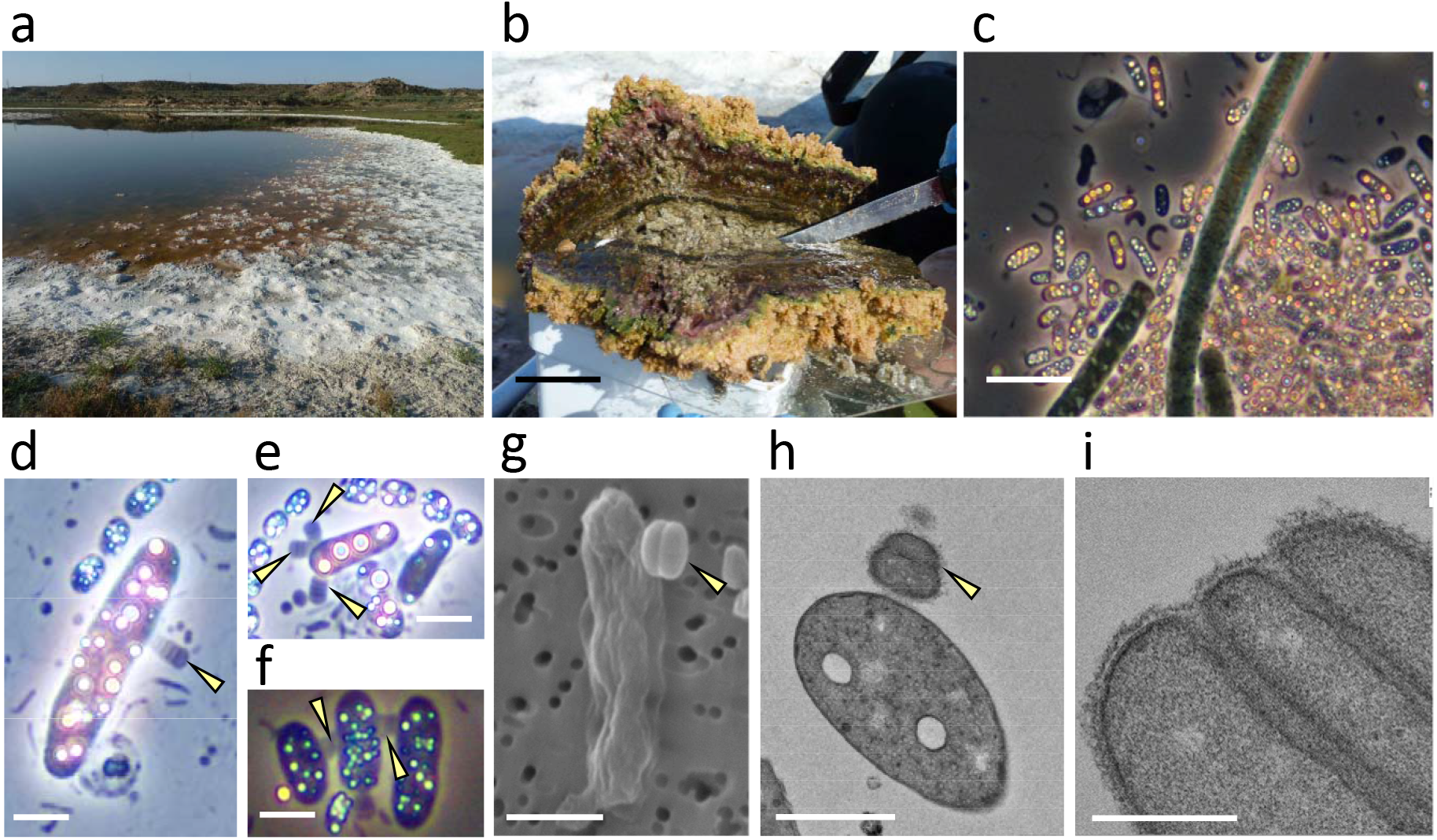
Sampling site and microscopy observation of *Vampirococcus* cells. **a**, General view of the microbial mat covering the shore of the Salada de Chiprana lake. **b**, Closer view of a microbial mat section. **c**, Natural population of blooming sulfide-dependent anoxygenic photosynthetic bacteria in waters of microbial mat containers after several weeks of growth in the laboratory; note the conspicuous refringent intracellular sulfur inclusions. **d-f**, Closer microscopy view of anoxygenic photosynthetic bacteria infected by epibiotic *Vampirococcus* cells and few-cell filaments (indicated by yellow arrows). **g**, Scanning electron microscopy image of a host cell infected by two stacking cells of *Vampirococcus* (yellow arrow). **h**, Transmission electron microscopy (TEM) image of a thin section of a host cell infected by *Vampirococcus* (yellow arrow). **i**, Closer TEM view of a thin section of *Vampirococcus* cells, notice the fibrous rugose cell surface and the large space separating contiguous cells. Scale bars: 5 cm (b), 5 μm (c), 1 μm (d-h), 0.5 μm (i).

Since the first *Vampirococcus* description included transmission electron microscopy (TEM) images, to further ascertain this identification we examined our Chiprana Lake samples under TEM and scanning electron microscopy (SEM). SEM images confirmed the peculiar structure of the epibionts, with multiple contiguous cells separated by deep grooves (Fig. 1g). Thin sections observed under TEM confirmed that the cells were actually separated by a space of ~20-50 nm filled by a fibrous material (Fig. 1h-i). The space between epibiont and host cells was larger (~100 nm) and also filled by dense fibrous material (Fig. 1h). The sections also showed that, in contrast with the typical Gram-negative double membrane structure of the host, the epibiont cells had a single membrane surrounded by a thick layer of fibrous material that conferred a rugose aspect to the cells (Fig. 1i). In sharp contrast with the often highly vacuolated cytoplasm of the host, the epibiont cells showed a dense, homogeneous content. These observations were also in agreement with those published for *Vampirococcus*, reinforcing our conclusion that the epibionts we observed belonged to this genus, although most likely to a different species, as the first *Vampirococcus* described occurred in a non-hypersaline lake^14^.

Using a micromanipulator coupled to an inverted microscope, we collected cells of the anoxygenic photosynthetic bacterium carrying *Vampirococcus* attached to their surface (Extended Data Fig. 1) and proceeded to amplify, clone, and sequence their 16S rRNA genes. We were able to obtain sequences for both the epibiont and the host for ten infected cells and, in all cases, we retrieved the same two sequences. The host was found to be a *Halochromatium*-like gammaproteobacterium (Extended Data Fig. 2). Phylogenetic analysis of the epibiont sequence showed that it branched within the CPR radiation close to the Absconditabacteria (Extended Data Fig. 3), previously known as candidate phylum SR1^2^. Since all host and epibiont cells we analyzed had identical 16S rRNA gene sequences, suggesting that they were the result of a clonal bloom, we collected three sets of ca. 10 infected cells and carried out whole genome amplification (WGA) before sequencing (Illumina HiSeq; see Methods). This strategy allowed us to assemble the nearly complete genome sequence of the *Vampirococcus* epibiont (see below). In contrast with the completeness of this genome, we only obtained a very partial assembly (~15%) of the host genome, probably because of the consumption of the host DNA by the epibiont. To make more robust phylogenetic analyses of *Vampirococcus*, we retrieved the protein sequence set used by Hug et al. to reconstruct a multi-marker large-scale phylogeny of bacteria^2^. The new multi-gene maximum likelihood (ML) phylogenetic tree confirmed the affiliation of our *Vampirococcus* species to the Absconditabacteria with maximum support, and further placed this clade within a larger well-supported group also containing the candidate phyla Gracilibacteria and Peregrinibacteria (Fig. 2a and Extended Data Fig. 4). Therefore, our epibiotic bacterium represents the first characterized member of this large CPR clade and provides a phylogenetic identity for the predatory bacterial genus *Vampirococcus* described several decades ago. We propose to call this new species *Candidatus* Vampirococcus lugosii (see Taxonomic Appendix).

**Fig. 2.**
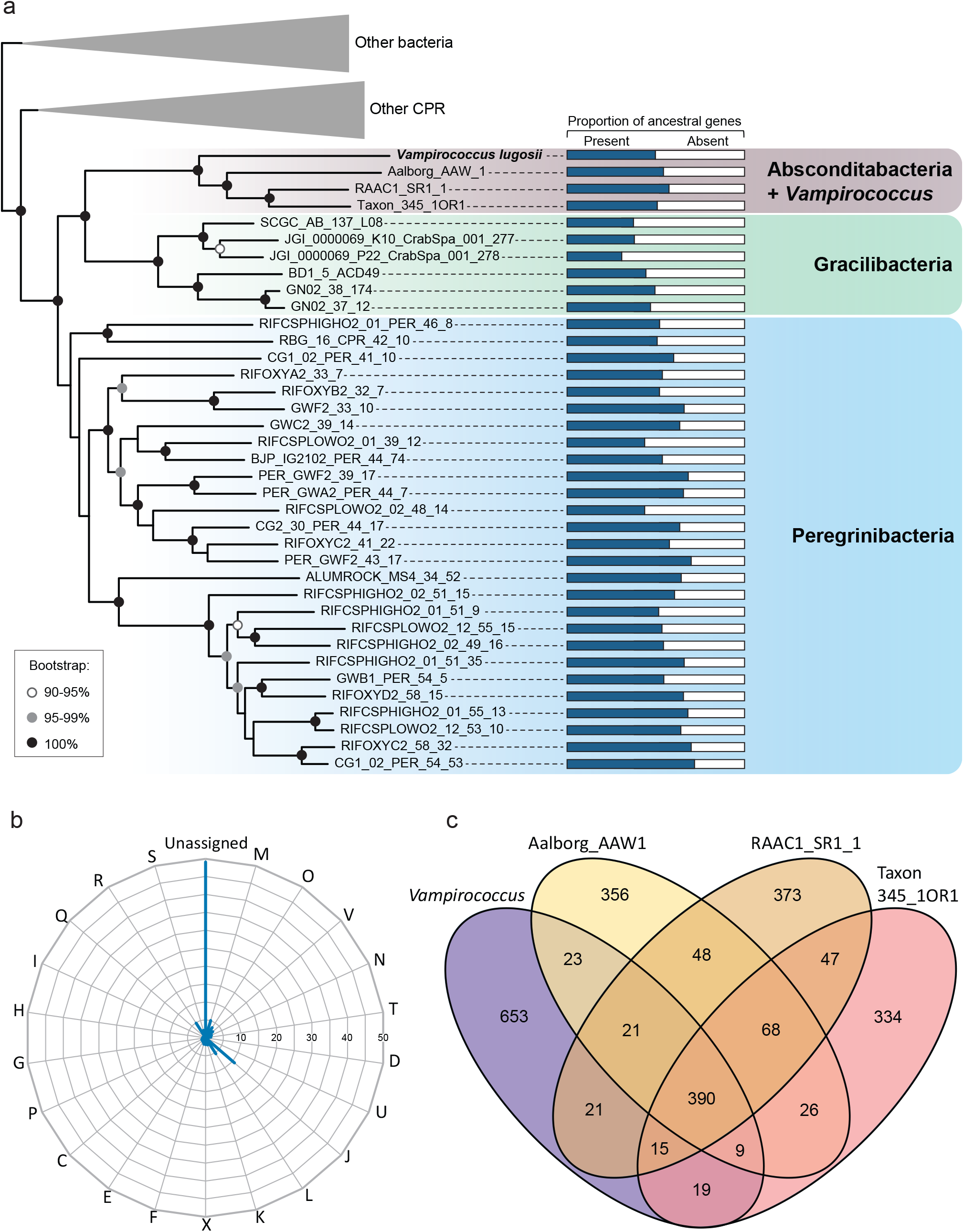
Phylogeny and global gene content of the *Vampirococcus* genome. **a**, Maximum likelihood phylogenetic tree of bacteria based on a concatenated dataset of 16 ribosomal proteins showing the position of *Vampirococcus lugosii* close to the Absconditabacteria (for the complete tree, see Extended Data Fig. 4). Histograms on the right show the proportion of genes retained in each species from the ancestral pool inferred for the last common ancestor of Absconditabacteria, Gracilibacteria and Peregrinibacteria. **b**, Percentage of *Vampirococcus* genes belonging to the different COG categories. **c**, Genes shared by *Vampirococcus* and the three Absconditabacteria genomes shown in the phylogenetic tree. COG categories are: Energy production and conversion [C]; Cell cycle control, cell division, chromosome partitioning [D]; Amino acid transport and metabolism [E]; Nucleotide transport and metabolism [F]; Carbohydrate transport and metabolism [G]; Coenzyme transport and metabolism [H]; Lipid transport and metabolism [I]; Translation, ribosomal structure and biogenesis [J]; Transcription [K]; Replication, recombination and repair [L]; Cell wall/membrane/envelope biogenesis [M]; Secretion, motility and chemotaxis [N]; Posttranslational modification, protein turnover, chaperones [O]; Inorganic ion transport and metabolism [P]; General function prediction only [R]; Function unknown [S]; Intracellular trafficking, secretion, and vesicular transport [U]; Defense mechanisms [V]; Mobilome: prophages, transposons [X]; Secondary metabolites biosynthesis, transport and catabolism [Q].

### Genomic evidence of adaptation to predatory lifestyle

We sequenced DNA from three WGA experiments corresponding each to ~10 *Halochromatium*-*Vampirococcus* consortia. Many of the resulting (57.2 Mb) raw sequences exhibited similarity to those of available Absconditabacteria/SR1 metagenome-assembled genomes (MAGs) and, as expected, some also to Gammaproteobacteria (host-derived sequences) as well as a small proportion of potential contaminants probably present in the original sample (*Bacillus*- and fungi-like sequences). To bin the *Vampirococcus* sequences out of this mini-metagenome, we applied tetranucleotide frequency analysis on the whole sequence dataset using emergent self-organizing maps (ESOM)^6^. One of the ESOM sequence bins was enriched in Absconditabacteria/SR1-like sequences and corresponded to the *Vampirococcus* sequences, which we extracted and assembled independently. This approach yielded an assembly of 1,310,663 bp. We evaluated its completeness by searching i) a dataset of 40 universally distributed single-copy genes^16^ and ii) a dataset of 43 single-copy genes widespread in CPR bacteria^8^. We found all them as single-copy genes in the *Vampirococcus* genome, with the exception of two signal recognition particle subunits from the first dataset which are absent in many other CPR bacteria^17^. These results supported that the *Vampirococcus* genome assembly was complete and had no redundancy. Manually curated annotation predicted 1,151 protein-coding genes, a single rRNA gene operon, and 38 tRNA coding genes. As already found in other Absconditabacteria/SR1 genomes^18^, the genetic code of *Vampirococcus* is modified, with the stop codon UGA reassigned as an additional glycine codon.

A very large proportion of the predicted proteins (48.95%) had no similarity to any COG class and lacked any conserved domain allowing their functional annotation (Fig. 2b). Thus, as for other CPR bacteria, a significant part of their cellular functions remains inaccessible. A comparison with three other Absconditabacteria genomes revealed a very small set of only 390 genes conserved in all them (Fig. 2c), suggesting a highly dynamic evolution of gene content in these species. Comparison with more distantly related CPR groups (Gracilibacteria and Peregrinibacteria) showed that gene loss has been a dominant trend in all these organisms, which have lost 30-50% of the ~1100 genes inferred to have existed in their last common ancestor (Fig. 2a). Nevertheless, this loss of ancestral genes was accompanied by the acquisition of new ones by different mechanisms, including horizontal gene transfer (HGT). In the case of *Vampirococcus*, we detected, by phylogenetic analysis of all individual genes that had homologs in other organisms, the acquisition of 126 genes by HGT from various donors (Extended Data Fig. 5).

The set of genes that could be annotated provided interesting clues about the biology and lifestyle of *Vampirococcus*. The most striking feature was its oversimplified energy and carbon metabolism map (Fig. 3). ATP generation in this CPR species appeared to depend entirely on substrate-level phosphorylation carried out by the phosphoenolpyruvate kinase (EC 2.7.1.40). In fact, *Vampirococcus* only possesses incomplete glycolysis, which starts with 3-phosphoglycerate as first substrate. This molecule is the major product of the enzyme RuBisCO and, therefore, most likely highly available to *Vampirococcus* from its photosynthetic host. Comparison with nearly complete MAG sequences available for other Absconditabacteria/SR1 showed that *Vampirococcus* has the most specialized carbon metabolism, with 3-phosphoglycerate as the only exploitable substrate, whereas the other species have a few additional enzymes that allow them to use other substrates (such as ribulose-1,5P and acetyl-CoA) as energy and reducing power (NADH) sources (Extended Data Fig. 6). This metabolic diversification probably reflects their adaptation to other types of hosts where these substrates are abundant. *Vampirococcus* also lacks all the enzymes involved in some Absconditabacteria/SR1 in the 3-phosphoglycerate-synthesizing AMP salvage pathway^19^, including the characteristic archaeal-like type II/III RuBisCO^20^.

**Fig. 3.**
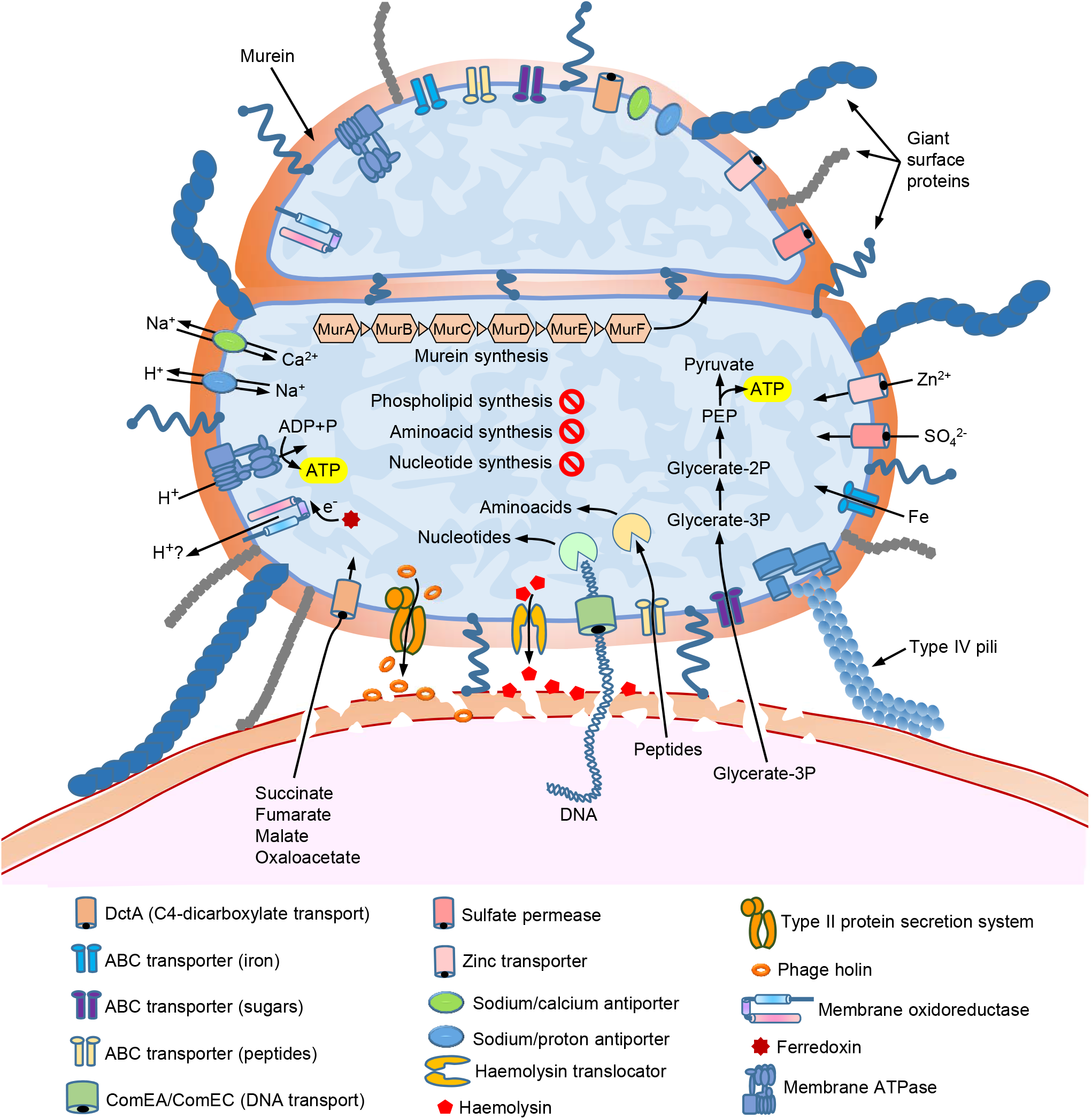
Metabolic and cell features inferred from the genes encoded in the *Vampirococcus* genome. The diagram shows the host cell surface (bottom) with two stacking *Vampirococcus* cells attached to its surface (as in Fig. 1h).

The genomes of Absconditabacteria/SR1 and *Vampirococcus* encode several electron carrier proteins (e.g., ferredoxin, cytochrome *b*_5_, several Fe-S cluster proteins) and a membrane F1FO-type ATP synthase. However, they apparently lack any standard electron transport chain and, therefore, they seem to be non-respiring^18,19,21^. The electron carrier proteins may be related with the oxidative stress response and/or the reoxidation of reduced ferredoxin or NADH^19^. In the absence of any obvious mechanism to generate proton motive force (PMF), the presence of the membrane ATP synthase is also intriguing. It has been speculated either that CPR bacteria might tightly adhere to their hosts and scavenge protons from them or that the membrane ATP synthase might work in the opposite direction as an ATPase, consuming ATP generated by substrate level phosphorylation to extrude protons and drive antiporters^9^. However, in the case of *Vampirococcus* the direct transport of protons from the host is unlikely since, as observed in the TEM sections (Fig. 1h), it seems that there is no direct contact with the host cell membrane. Parasite and host cell membranes are separated by a relatively large space of ~100 nm, which would be largely conducive to proton diffusion and inefficient transfer between cells. Although the proton/cation antiporters (e.g., for Na^+^, K^+^ or Ca^+^) encoded by *Vampirococcus* and the other Absconditabacteria/SR1 may serve to produce some PMF, it is improbable that this mechanism represents a major energy transducing system as cells would accumulate cations and disrupt their ionic balance; these antiporters are most likely involved in cation homeostasis. This prompted us to investigate other ways that these cells might use to generate PMF usable by their ATP synthase. We found a protein (Vamp_33_45) with an atypical tripartite domain structure. The N-terminal region, containing 8 transmembrane helices, showed similarity with several flavocytochromes capable of moving electrons and/or protons across the plasma membrane (e.g.,^22^). The central part of the protein was a rubredoxin-like nonheme iron-binding domain likely able to transport electrons. Finally, the C-terminal region, containing an NAD-binding motif, was similar to ferredoxin reductases involved in electron transfer^23^. This unusual *Vampirococcus* 3-domain protein is well conserved in the other Absconditabacteria/SR1 genomes sequenced so far, suggesting it plays an important function in this CPR phylum. Its architecture suggests that it can transport electrons and/or protons across the membrane using ferredoxin as electron donor and makes it a strong candidate to participate in a putative new PMF-generating system. Alternatively, this protein could play a similar role to that of some oxidoreductases in the strict anaerobic archaeon *Thermococcus onnurineus*, including a thioredoxin reductase, which couple reactive oxygen species detoxification with NAD(P)+ regeneration from NAD(P)H to maintain the intracellular redox balance and enhance O_2_-mediated growth despite the absence of heme-based or cytochrome-type proteins^24^.

Despite our *Vampirococcus* genome sequence appears to be complete, genes encoding enzymes involved in the biosynthesis of essential cell building blocks such as amino acids, nucleotides and nucleosides, cofactors, vitamins, and lipids are almost completely absent (Fig. 2b). Therefore, the classical bacterial metabolic pathways for their synthesis^25^ do not operate in *Vampirococcus*. Such extremely simplified metabolic potential, comparable to that of intracellular parasitic bacteria such as *Mycoplasma*^26^, implies that *Vampirococcus* must acquire these molecules from an external source and supports the predatory nature of the interaction with its photosynthetic host. An intriguing aspect of this interaction concerns the transfer of substrates from the host to *Vampirococcus*, especially considering that, despite examination of several serial ultrathin sections, the cell membranes of these two partners do not appear to be in direct contact (Fig. 1h). *Vampirococcus* encodes several virulence factors, including divergent forms of hemolysin and hemolysin translocator (Vamp_11_169 and Vamp_9_166, respectively), a phage holin (Vamp_5_129), and a membrane-bound lytic murein transglycosylase (Vamp_144_2). These proteins are likely involved in the host cell wall and membrane disruption leading to cell content release. Hemolysin has also been found in Saccharibacteria (formerly candidate phylum TM7), the only CPR phylum for which an epibiotic parasitic lifestyle has been demonstrated so far^12,13^. Recent coupled lipidomic-metagenomic analyses have shown that CPR bacteria that lack complete lipid biosynthesis are able to recycle membrane lipid from other bacteria^27^. In *Vampirococcus*, also devoid of phospholipid synthesis, a phospholipase gene (Vamp_34_196) predicted to be secreted and that has homologs involved in host phospholipid degradation in several parasitic bacteria^28^, may not only help disrupting the host membrane but also to generate a local source of host phospholipids that it can use to build its own cell membrane. Two *Vampirococcus* peptidoglycan hydrolases (Vamp_68-56_103 and Vamp_145_30) and one murein DD-endopeptidase (Vamp_311_38), also predicted to be secreted, most probably contribute to degrade the host cell wall.

*Vampirococcus* also possesses a number of genes encoding transporters, most of them involved in the transport of inorganic molecules (Fig. 3). One notable exception is the competence-related integral membrane protein ComEC (Vamp_67_106)^29^ which, together with ComEA (Vamp_21_186) and type IV pili (see below), probably plays a role in the uptake of host DNA that, once transported into the epibiont, can be degraded by various restriction endonucleases (five genes encoding them are present) and recycled to provide the nucleotides necessary for growth (Extended Data Fig. 7). These proteins are widespread in other CPR bacteria where they may have a similar function^30^. *Vampirococcus* also encodes an ABC-type oligopeptide transporter (Vamp_40_40) and a DctA-like C4-dicarboxylate transporter (Vamp_41_97), known to catalyze proton-coupled symport of several Krebs cycle dicarboxylates (succinate, fumarate, malate, and oxaloacetate)^31^. The first, coupled with the numerous peptidases present in *Vampirococcus*, most likely is a source of amino acids. By contrast, the role of DctA is unclear since *Vampirococcus* does not have a Krebs cycle.

In sharp contrast with its simplified central metabolism, *Vampirococcus* possesses genes related to the construction of an elaborate cell surface, which seems to be a common theme in many CPR bacteria^9,11^. They include genes involved in peptidoglycan synthesis, several glycosyltransferases, a Sec secretion system, and a rich repertoire of type IV pilus proteins. The retractable type IV pili are presumably involved in the tight attachment of *Vampirococcus* to its host and in DNA uptake in cooperation with the ComEC protein. Other proteins probably play a role in the specific recognition and fixation to the host, including several very large proteins. In fact, the *Vampirococcus* membrane proteome is enriched in giant proteins. The ten longest predicted proteins (between 1392 and 4163 aa, see Extended Data Fig. 8) are inferred to have a membrane localization and are probably responsible of the conspicuous fibrous aspect of its cell surface (Fig. 1i). Most of these proteins possess domains known to be involved in the interaction with other molecules, including protein-protein (WD40, TRP, and PKD domains) and protein-lipid (saposin domain) interactions and cell adhesion (DUF11, integrin, and fibronectin domains). Two other large membrane proteins (Vamp_6_203, 2368 aa, and Vamp_19_245, 1895 aa) may play a defensive role as they contain alpha-2-macroglobulin protease-inhibiting domains that can protect against proteases released by the host. Several other smaller proteins complete the membrane proteome of *Vampirococcus*, some of them also likely involved in recognition and attachment to the host thanks to a variety of protein domains, such as VWA (Vamp_41_85) and flotillin (Vamp_11_100).

### New CRISPR-Cas systems and other defense mechanisms in *Vampirococcus*

Although most CPR phyla are devoid of CRISPR-Cas^32^, some have been found to contain new systems with original effector enzymes such as CasY^33^. In contrast with most available Absconditabacteria genomes, *Vampirococcus* possesses two CRISPR-Cas loci (Fig. 4a and Extended Data Fig. 9). The first is a class II type V system that contains genes coding for Cas1, Cas2, Cas4, and Cpf1 proteins associated to 34 spacer sequences of 26-32 bp. Proteins similar to those of this system are encoded not only in genomes of close relatives of the Absconditabacteria (Gracilibacteria and Peregrinibacteria) but in many other CPR phyla. These sequences form monophyletic groups in phylogenetic analyses (e.g., Cas1, see Fig. 4b), which suggests that this type V system is probably ancestral in these CPR. The second system found in *Vampirococcus* belongs to the class I type III and contains genes coding for Cas1, Cas2, Csm3, and Cas10/Csm1 proteins associated to a cluster of 20 longer (35-46 bp) spacers. In contrast with the previous CPR-like system, the proteins of this second system did not show strong similarity with any CPR homologue but with sequences from other bacterial phyla, suggesting that they have been acquired by HGT. Phylogenetic analysis confirmed this and supported that *Vampirococcus* gained this CRISPR-Cas system from different distant bacterial donors (Extended Data Fig. 10). Interestingly, these two CRISPR-Cas systems encode a number of proteins that may represent new effectors. A clear candidate is the large protein Vamp_48_93 (1158 aa), located between Cpf1 and Cas1 in the type V system (Fig. 4a), which contains a DNA polymerase III PolC motif. Very similar sequences can be found in a few other CPR (some Roizmanbacteria, Gracilibacteria, and Portnoybacteria) and in some unrelated bacteria (Extended Data Fig. 11). As in *Vampirococcus*, the gene coding for this protein is contiguous to genes encoding different Cas proteins in several of these bacteria, including Roizmanbacteria, Omnitrophica, and the deltaproteobacterium *Smithella* sp. (Extended Data Fig. 11). This gene association, as well as the very distant similarity between this protein and Cpf1 CRISPR-associated proteins of bacterial type V systems, supports that it is a new effector in type V CRISPR-Cas systems. Additional putative new CRISPR-associated proteins likely exist also in the *Vampirococcus* type III system (Fig. 4a). Three proteins encoded by contiguous genes (Vamp_21_116, Vamp_21_127, and Vamp_21_128) exhibit very distant similarity with type III-A CRISPR-associated RAMP (Repeat Associated Mysterious Proteins) Csm4, Csm5, and Csm6 sequences, respectively, and most probably represent new RAMP subfamilies. To date, Absconditabacteria^34^ and Saccharibacteria^35^ are the only CPR phyla for which phages have been identified. Because of its proximity to Absconditabacteria, *Vampirococcus* is probably infected by similar phages, so that the function of its CRISPR-Cas systems may be related to the protection against these genetic parasites. Nevertheless, we did not find any similarity between the *Vampirococcus* spacers and known phage sequences, suggesting that it is infected by unknown phages. Alternatively, considering that *Vampirococcus* -as most likely many other CPR bacteria-seems to obtain nucleotides required for growth by uptaking host DNA, an appealing possibility is that the CRISPR-Cas systems participate in the degradation of the imported host DNA.

**Fig. 4.**
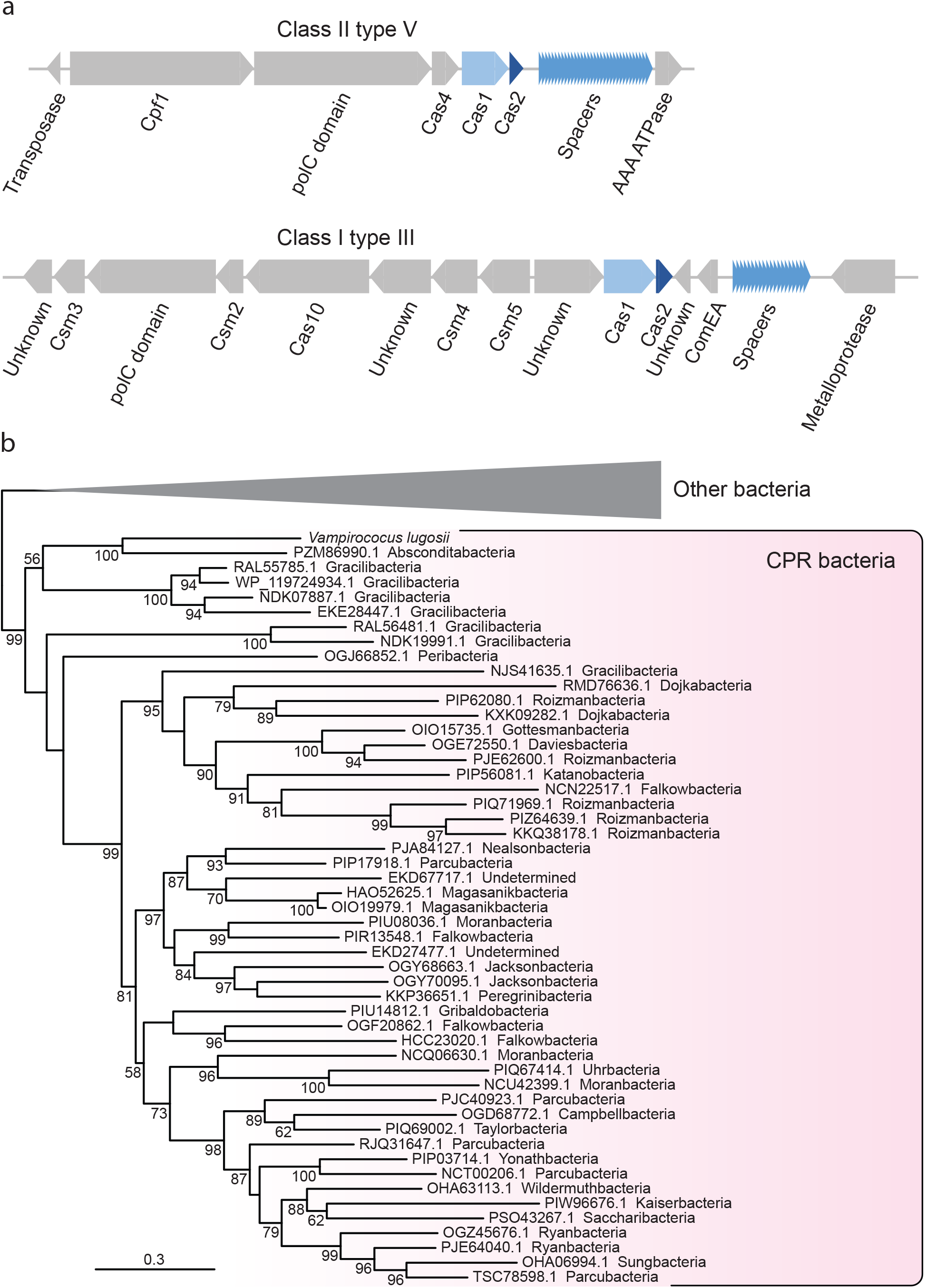
CRISPR-Cas systems in *Vampirococcus*. **a**, Genes in the two systems encoded in the *Vampirococcus lugosii* genome. **b**, Maximum likelihood phylogenetic tree of the Cas1 protein encoded in the class II type V system, numbers at branches indicate bootstrap support.

Although CPR bacteria have been hypothesized to be largely depleted of classical defense mechanisms^36^, we found that *Vampirococcus*, in addition to the two CRISP-Cas loci, is endowed with various other protection mechanisms. These include an AbiEii-AbiEi Type IV toxin-antitoxin system, also present in other CPR bacteria, which may offer additional protection against phage infection^37^ and several restriction-modification systems, with three type I, one type II and one type III restriction enzymes and eight DNA methylases. In addition to a defensive role, these enzymes may also participate in the degradation of the host DNA. As in its sister-groups Absconditabacteria and Gracilibacteria^5,18,19,38,39^, *Vampirococcus* has repurposed the UGA stop codon to code for glycine. The primary function of this recoding remains unknown but it has been speculated that it creates a genetic incompatibility, whereby these bacteria would be “evolutionarily isolated" from their environmental neighbors, preventing their potential competitors from acquiring their genomic innovations by HGT^18^. However, the opposite might be argued as well, since the UGA codon reassignment can protect *Vampirococcus* from foreign DNA expression upon uptake by leading to aberrant protein synthesis via read-through of the UGA stop with Gly insertion. This can be important for these CPR bacteria because they are not only impacted by phages^34^ but they most likely depend on host DNA import and degradation to fulfill their nucleotide requirements. In that sense, it is interesting to note that the *Vampirococcus* ComEA protein likely involved in DNA transport^40^ is encoded within the class I type III CRISPR-Cas system (Fig. 4a).

### Concluding remarks

Our phylogenomic analyses show that *Vampirococcus lugosii*, the first genomically characterized example of the rare “predatory” bacteria described several decades ago^14^, is closely related to members of the phylum Absconditabacteria. *V. lugosii* thus becomes the first species characterized for a large subgroup of CPR bacteria that also includes Gracilibacteria and Peregrinibacteria (Fig. 2a). *Vampirococcus* is a parasite that attaches to the surface of anoxygenic photosynthetic gammaproteobacteria and kills its host by consumption of its cytoplasmic content. In agreement with this predatory nature, the *Vampirococcus* genome sequence codes for an extremely simplified metabolism, implying that it depends on its host to obtain all cell constituents (amino acids, lipids, nucleotides, nucleosides, etc.). By contrast, it codes for a complex cell surface, including peptidoglycan, many transporters, pili, and several giant proteins likely involved in the recognition and attachment to the host. The sticky nature of the *Vampirococcus* cell surface seems crucial for an original transmission mode. As *Vampirococcus* cells divide, they form stalks of piling cells, therefore increasing the probability of apical cells to encounter and attach to another host cell. Parasite filaments can connect two host cells but also release apical cells that, upon attachment to the host, start dividing again. Despite intensive observations of TEM thin sections, communication bridges between parasite and host, and between piling parasite cells, were never observed. This implies a unique feeding mode whereby the parasite partially degrades the host cell membrane, exploiting its membrane hydrolysis products and cytoplasmic content. In principle, this feeding mode would be only available for the parasite in immediate vicinity with the host cell. This would entail that, upon cell division, stalk parasite cells would be inactive until they stick to another host cell. Alternatively, although more speculative, stalk parasite cells might depend on the same original host and feed from neighboring parasitic cells by some sort of facilitated diffusion through the fibrous cell envelope in a peculiar type of kin-feeding mode. Despite a clear trend towards metabolic simplification, *Vampirococcus*, as well as other related CPR bacteria, presents a high degree of evolutionary innovation. It includes many genes that do not show any similarity to those in sequence databases and that lack known conserved domains. Nevertheless, in some cases the genomic context allows to propose possible functions for some of these genes, as in the case of new effector proteins associated to the CRISPR-Cas loci. Other interesting innovations have evolved through new unique combinations of protein domains, as in the case of the tripartite domain protein putatively involved in the generation of a proton gradient across membrane trough an unconventional respiratory chain.

So far, only members of the CPR Saccharibacteria have been maintained in stable cultures, allowing the characterization of their parasitic relationship with various bacterial hosts^12,13,41^. Using a “reverse genomics” approach with specific antibodies and cell sorting with flow cytometry, cultures of human oral Absconditabacteria/SR1 species mixed with other bacteria have been recently established but their poor growth rate did not enable the detailed characterization of their biology^13^. Although we did not achieve to get stable *Vampirococcus* cultures, the recurrent examination of living material with optical microscopy combined with electron microscopy observations and genome sequence analysis clearly demonstrate its predatory nature and allowed us to infer several mechanisms for host exploitation. Similarities in gene content with relatives in the Absconditabacteria/SR1 group suggest that this whole clade is composed of predatory or parasitic species. They most likely infect a wide spectrum of bacterial hosts, both photosynthetic and non-photosynthetic, as deduced from their metabolic specializations and their occurrence in very different habitats, such as deep aquifers, lakes, and the human body^13,18,19^. The same has been proved for the Saccharibacteria^12,13^, adding to the growing evidence for parasitism as the general lifestyle of CPR bacteria. The vast diversity and habitat distribution of these bacteria suggest that they may play an ecological role analogous to that of viruses by controlling the population size of their host bacteria.

### Taxonomic Appendix

The genome sequence of a species labelled as “*Candidatus* Vampirococcus archaeovorus” already existed in GenBank (CP019384.1) and belonged to a parasite of methanogenic archaea ascribed to the candidate division OP3, now known as Omnitrophica^42^. However, the organism that we have studied in this work fits much better the original description of the genus *Vampirococcus* as an epibiotic parasite of photosynthetic anoxygenic bacteria with flat stacked cells^14^. Therefore, because of historical priority reasons, the genus name *Vampirococcus*, from "vampire" (Serbian: vampir, blood-sucker) and "coccus" (Greek: coccus, grain or berry), must be retained for this organism and a different genus name needs to be proposed for the homonymous parasite of archaea affiliated to the Omnitrophica.

### Description of ‘ *Candidatus* Vampirococcus lugosii’

Lugosii after Bela Lugosi (1882-1956), who played the role of the vampire in the iconic 1931’s film “Dracula”. Epibiotic bacterium that preys on anoxygenic photosynthetic gammaproteobacterial species of the genus *Halochromatium*. Non-flagellated, small flat rounded cells (500-600 nm diameter and 200-250 nm height) that form piles of up to 10 cells attached to the surface of the host. Gram-positive cell wall structure. Complete genome sequence, GenBank/EMBL/DDBJ accession number XXXX.

## Methods

### Sample collection and processing

Microbial mats were sampled in December 2013 in the Salada de Chiprana (NE Spain) permanent athalassic hypersaline lake (55.9 g total dissolved salt l^−1^, pH 8.23) at a depth of ~50 cm. Fragments of these microbial mats were maintained under artificial illumination in 20 l plastic containers filled with filtered lake water (<0.2 μm). We daily monitored the growth of planktonic bacteria in these containers by optical microscopy. After ~3 weeks, we observed a bloom of *Chromatium*-like anoxygenic photosynthetic bacteria and noticed that many cells had smaller epibiotic cells attached to them. We collected individual consortia of host-epibiont cells using an Eppendorf PatchMan NP2 micromanipulator equipped with 6 μm-diameter microcapillaries (Eppendorf) mounted on a Leica Dlll3000 B inverted microscope. Consortia were rinsed twice with sterile 10 mM Tris pH 8.0 buffer and finally deposited in a volume of 0.5 μl of this buffer and stored frozen at −20°C until further processing.

### Microscopy

Light microscopy observation of living material was carried out on a Zeiss Axioplan microscope equipped with a Nikon Coolpix B500 color camera. For Scanning Electron Microscopy (SEM), liquid samples enriched in *Vampirococcus* and its host were deposited on top of 0.1 μm pore-diameter filters (Whatman) under a mild vacuum aspiration regime and briefly rinsed with 0.1-μm filtered and autoclaved MilliQ water under the same vacuum regime.

Filters were let dry and sputtered with gold. SEM observations were carried out using a Field Emission Electron Microscope Philips XL30 S-FEG. Secondary electron (SE2) images were acquired using an In Lens detector at an accelerating voltage of 2.0 kV and a working distance of ∼7.5 mm. For Transmission Electron Microscopy (TEM), 1.5 ml of the sample were centrifuged for 2 minutes at 10,000 rpm. Cells were fixed with 2% glutaraldehyde in 0.1 M cacodylate at pH 7.4 and postfixed in 2% osmium tetroxide for 30 minutes. Cells were then dehydrated in a gradual series of ethanol baths (50%, 70% and 100%) and dried cells were embedded in epoxy resin. The resin block was cut into 90 nm thick sheets using an Ultracut UCT ultramicrotome. The sheets were placed on copper grids and observed in a JEM1010 (Jeol) microscope with an acceleration voltage of 100 KeV.

### DNA extraction, 16S rRNA gene PCR amplification, and whole genome amplification

DNA was purified from individual (or few, ca. 10 cells) cell consortia with the PicoPure DNA extraction kit (Applied Biosystems). 16S rRNA genes were PCR-amplified using the primers B-27F (AGAGTTTGATCCTGGCTCAG) and 1492R (GGTTACCTTGTTACGACTT). PCR reactions were performed for 30 cycles (denaturation at 94°C for 15 s, annealing at 55°C for 30 s, extension at 72°C for 2 min) preceded by 2 min denaturation at 94°C, and followed by 7 min extension at 72°C. 16S rRNA gene clone libraries were constructed with the Topo TA cloning system (Invitrogen) following the instructions provided by the manufacturer. After plating, positive transformants were screened by PCR amplification using M13R and T7 flanking vector primers. 16S rRNA amplicons were Sanger-sequenced using the 1492R primer by Genewiz (Essex, UK). Whole genome amplification (WGA) was carried out on PicoPure-extracted DNA using Multiple Displacement Amplification^43^ (MDA) with the REPLI-g WGA kit (Qiagen) and Multiple Annealing and Looping-Based Amplification Cycles^44^ (MALBAC) with the MALBAC Single Cell WGA kit (Yikon Genomics).

### Genome sequencing, assembly and annotation

Sequencing of WGA products from three independent consortia with identical 16S rRNA gene sequences (V7 and V8, amplified by MDA, and V12, amplified by MALBAC) was done using Illumina HiSeq2500 v4 (2×125 bp paired-end reads) by Eurofins Genomics (Ebersberg, Germany). Each dataset was separately assembled into contigs with SPAdes v3.6.0^45^ using default parameters. MDA contigs longer than 2.5 kb were clustered using the ESOM procedure (ESOM Tools v. 1.1;^46^) along with 3 reference genomes chosen for their close phylogenetic relation to *Vampirococcus* and its host: Candidate division SR1 bacterium RAAC1_SR1_1 (CP006913), *Allochromatium vinosum* DSM 180 (NC_013851), *Thioflavicoccus mobilis* 8321 (NC_019940), along with an external reference: *Escherichia coli* BL21-Gold(DE3)pLysS AG (NC_012947). The clustering was done on nucleotide tetramer frequency distributions computed with the tetramer_freqs_esom.pl script^6^ using a window size of 5 kb. ESOM training parameters where set to k-batch training method, 140 x 250 mesh size, radius start 50, 20 training epochs. All other parameters were set to their defaults. The visualization of the ESOM maps was done with a U-Matrix background and inverse gray-scale gradient coloring (Extended Data Fig. 12). Further improvement was achieved by cross-matching the MDA ESOM-processed sequence set with contigs from the MALBAC set, which increased the Vampirococcus total sequence size by ~10%. The completeness and contamination of the bins were assessed using CheckM^47^.

CDS prediction was performed on the assembled contigs using Prodigal version 2.6.2 (single mode, translation table 25 (Candidate Division SR1 and *Gracilibacteria*)^48^. For more accurate functional annotation, we submitted the amino acid sequence of predicted genes to the blastp command of DIAMOND version 0.7.9 (maximum e-value of 10-5)^49^ to search three databases: the non-redundant protein RefSeq (release 68; Nov 3, 2014), COG^50^, and SEED (release of September 14, 2011)^51^. Enzymes involved in metabolic pathways were also searched against the KEGG database^25^. Conserved protein motifs were searched using SMART^52^. Prediction of transmembrane helices was done with the TMHMM Server v. 2.0 (http://www.cbs.dtu.dk/services/TMHMM/). CRISPR-Cas loci were identified using CRISPRCasFinder^53^ and CRISPRminer^54^. Ancestral gene content and gene loss in the group Absconditabacteria + Gracilibacteria + Peregrinibacteria were inferred by first carrying out orthologue clustering with OrthoFinder v1.1.20^55^ with default parameters, and then applying the Dollo parsimony method implemented in the Count software^56^.

### Phylogenetic Analyses

16S rRNA and protein sequences were aligned using MAFFT L-INS-i^57^ and poorly aligned regions were removed with trimAl –automated1^58^. Maximum Likelihood (ML) phylogenetic trees were reconstructed using IQ-tree v. 1.5.549^59^ with the GTR+G+I model for 16S rRNA gene alignments and the LG+C20+G model for individual protein alignments. Multi-protein datasets were constructed by concatenation of individual protein trimmed alignments using SequenceMatrix^60^. ML trees of multi-protein datasets were inferred using IQ-tree and the LG+PMSF(C60)+F+G4 model with a guide tree inferred with the LG+C60+F+G4 model. In all cases, branch support was estimated with the bootstrap method (100 replicates) implemented in IQ-tree.

## Acknowledgements

We thank M. Ragon and the UNICELL platform (http://www.deemteam.fr/en/unicell) for carrying out WGA experiments, and A. Saghaï for 16S rRNA gene amplification assays of *Vampirococcus* relatives. This work was supported by the European Research Council (ERC) Advanced Grants ‘Protistworld’ and ‘Plast-Evol’ (322669 and 787904, respectively).

## Author contributions

D.M., P.L.-G., A.I.L.-A. and M.I. carried out sampling and monitored *Vampirococcus* growth. A.I.L.-A. carried out electron microscopy observations. D.M. micromanipulated cells. P.L.-G. amplified 16S rRNA genes from isolated cells. Y.Z. assembled and annotated the *Vampirococcus* genome. D.M. analyzed gene sequences to infer metabolism and other cell features. D.M. wrote the manuscript with help from the rest of co-authors.

## Notes

### Competing Interest Statement

The authors have declared no competing interest.

## References

1. López-García, P. & Moreira, D. Tracking microbial biodiversity through molecular and genomic ecology. Res. Microbiol. 159, 67–73 (2008).

2. Hug, L.A. et al. A new view of the tree of life. Nat. Microbiol. 1, 16048 (2016).

3. Solden, L., Lloyd, K. & Wrighton, K. The bright side of microbial dark matter: lessons learned from the uncultivated majority. Curr. Opin. Microbiol. 31, 217–226 (2016).

4. Castelle, C.J. & Banfield, J.F. Major new microbial groups expand diversity and alter our understanding of the Tree of Life. Cell 172, 1181–1197 (2018).

5. Rinke, C. et al. Insights into the phylogeny and coding potential of microbial dark matter. Nature 499, 431–437 (2013).

6. Dick, G.J. et al. Community-wide analysis of microbial genome sequence signatures. Genome Biol. 10, R85 (2009).

7. Zaremba-Niedzwiedzka, K. et al. Asgard archaea illuminate the origin of eukaryotic cellular complexity. Nature 541, 353–358 (2017).

8. Brown, C.T. et al. Unusual biology across a group comprising more than 15% of domain Bacteria. Nature 523, 208–211 (2015).

9. Castelle, C.J. et al. Biosynthetic capacity, metabolic variety and unusual biology in the CPR and DPANN radiations. Nat. Rev. Microbiol. 16, 629–645 (2018).

10. Dombrowski, N., Lee, J.H., Williams, T.A., Offre, P. & Spang, A. Genomic diversity, lifestyles and evolutionary origins of DPANN archaea. FEMS Microbiol. Lett. 366, fnz008 (2019).

11. Luef, B. et al. Diverse uncultivated ultra-small bacterial cells in groundwater. Nat. Commun. 6, 6372 (2015).

12. He, X. et al. Cultivation of a human-associated TM7 phylotype reveals a reduced genome and epibiotic parasitic lifestyle. Proc. Natl. Acad. Sci. U. S. A. 112, 244–249 (2015).

13. Cross, K.L. et al. Targeted isolation and cultivation of uncultivated bacteria by reverse genomics. Nat. Biotechnol. 37, 1314–1321 (2019).

14. Guerrero, R. et al. Predatory prokaryotes: predation and primary consumption evolved in bacteria. Proc. Natl. Acad. Sci. U. S. A. 83, 2138–42. (1986).

15. De Wit, R. Lake La Salada de Chiprana (NE Spain), an example of an athalassic salt lake in a cultural landscape. in Lake Sciences and Climate Change (ed. Nageeb-Rashed, M.) 43–60 (Intech ed., London, 2016).

16. Creevey, C.J., Doerks, T., Fitzpatrick, D.A., Raes, J. & Bork, P. Universally distributed single-copy genes indicate a constant rate of horizontal transfer. PLoS One 6, e22099 (2011).

17. Nelson, W.C. & Stegen, J.C. The reduced genomes of Parcubacteria (OD1) contain signatures of a symbiotic lifestyle. Front. Microbiol. 6, 713 (2015).

18. Campbell, J.H. et al. UGA is an additional glycine codon in uncultured SR1 bacteria from the human microbiota. Proc. Natl. Acad. Sci. U. S. A. 110, 5540–5545 (2013).

19. Kantor, R.S. et al. Small genomes and sparse metabolisms of sediment-associated bacteria from four candidate phyla. MBio 4, 00708–00713 (2013).

20. Sato, T., Atomi, H. & Imanaka, T. Archaeal type III RuBisCOs function in a pathway for AMP metabolism. Science. 315, 1003–1006. (2007).

21. Dueholm, M.S. et al. Complete genome sequence of the bacterium Aalborg_AAW-1, representing a novel family within the Candidate Phylum SR1. Genome Announc. 3, e00624–15 (2015).

22. Loschi, L. et al. Structural and biochemical identification of a novel bacterial oxidoreductase. J. Biol. Chem. 279, 50391–50400. (2004).

23. Arakaki, A.K., Ceccarelli, E.A. & Carrillo, N. Plant-type ferredoxin-NADP+ reductases: a basal structural framework and a multiplicity of functions. Faseb J. 11, 133–140 (1997).

24. Lee, S.H., Youn, H., Kang, S.G. & Lee, H.S. Oxygen-mediated growth enhancement of an obligate anaerobic archaeon *Thermococcus onnurineus* NA1. J. Microbiol. 57, 138–142 (2019).

25. Kanehisa, M. Enzyme annotation and metabolic reconstruction using KEGG. Methods Mol. Biol. 1611, 135–145 (2017).

26. Fraser, C.M. et al. The minimal gene complement of *Mycoplasma genitalium*. Science. 270, 397–403. (1995).

27. Probst, A.J. et al. Lipid analysis of CO(2)-rich subsurface aquifers suggests an autotrophy-based deep biosphere with lysolipids enriched in CPR bacteria. ISME J. 14, 1547–1560 (2020).

28. Toledo, A. & Benach, J.L. Hijacking and use of host lipids by intracellular pathogens. Microbiol. Spectr. 3, 0001–2014 (2015).

29. Hahn, J., Inamine, G., Kozlov, Y. & Dubnau, D. Characterization of comE, a late competence operon of *Bacillus subtilis* required for the binding and uptake of transforming DNA. Mol. Microbiol. 10, 99–111. (1993).

30. Meheust, R., Burstein, D., Castelle, C.J. & Banfield, J.F. The distinction of CPR bacteria from other bacteria based on protein family content. Nat. Commun. 10, 4173 (2019).

31. Groeneveld, M., Weme, R.G., Duurkens, R.H. & Slotboom, D.J. Biochemical characterization of the C4-dicarboxylate transporter DctA from *Bacillus subtilis*. J. Bacteriol. 192, 2900–2907 (2010).

32. Burstein, D. et al. Major bacterial lineages are essentially devoid of CRISPR-Cas viral defence systems. Nat. Commun. 7, 10613 (2016).

33. Chen, L.X. et al. Candidate Phyla Radiation Roizmanbacteria from hot springs have novel and unexpectedly abundant CRISPR-Cas systems. Front. Microbiol. 10, 928 (2019).

34. Paez-Espino, D. et al. Uncovering Earth’s virome. Nature 536, 425–430 (2016).

35. Dudek, N.K. et al. Novel microbial diversity and functional potential in the marine mammal oral microbiome. Curr. Biol. 27, 3752–3762 (2017).

36. Koonin, E.V., Makarova, K.S. & Wolf, Y.I. Evolutionary genomics of defense systems in Archaea and Bacteria. Annu. Rev. Microbiol. 71, 233–261 (2017).

37. Dy, R.L., Przybilski, R., Semeijn, K., Salmond, G.P. & Fineran, P.C. A widespread bacteriophage abortive infection system functions through a Type IV toxin-antitoxin mechanism. Nucleic Acids Res. 42, 4590–4605 (2014).

38. Wrighton, K.C. et al. Fermentation, hydrogen, and sulfur metabolism in multiple uncultivated bacterial phyla. Science 337, 1661–1665 (2012).

39. Hanke, A. et al. Recoding of the stop codon UGA to glycine by a BD1-5/SN-2 bacterium and niche partitioning between Alpha- and Gammaproteobacteria in a tidal sediment microbial community naturally selected in a laboratory chemostat. Front. Microbiol. 5, 231 (2014).

40. Seitz, P. et al. ComEA is essential for the transfer of external DNA into the periplasm in naturally transformable *Vibrio cholerae* cells. PLoS Genet. 10, e1004066 (2014).

41. Bor, B. et al. Insights obtained by culturing Saccharibacteria with their bacterial hosts. J. Dent. Res. 99, 685–694 (2020).

42. Kizina, J. et al. Insights into the biology of Candidate Division OP3 LiM populations. PhD thesis. Bremen, Germany, University of Bremen, 85–160 (2017).

43. Dean, F.B. et al. Comprehensive human genome amplification using multiple displacement amplification. Proc. Natl. Acad. Sci. U. S. A. 99, 5261–5266 (2002).

44. Zong, C., Lu, S., Chapman, A.R. & Xie, X.S. Genome-wide detection of single-nucleotide and copy-number variations of a single human cell. Science 338, 1622–1626 (2012).

45. Bankevich, A. et al. SPAdes: a new genome assembly algorithm and its applications to single-cell sequencing. J. Comput. Biol. 19, 455–477 (2012).

46. Ultsch, A. & Mörchen, F. ESOM-Maps: tools for clustering, visualization, and classification with Emergent SOM. (Tech. Rep. Dept. Math. Comput. Sci. Univ. Marburg, Ger., 2005).

47. Parks, D.H., Imelfort, M., Skennerton, C.T., Hugenholtz, P. & Tyson, G.W. CheckM: assessing the quality of microbial genomes recovered from isolates, single cells, and metagenomes. Genome Res. 25, 1043–1055 (2015).

48. Hyatt, D. et al. Prodigal: prokaryotic gene recognition and translation initiation site identification. BMC Bioinformatics 11, 1471–2105 (2010).

49. Buchfink, B., Xie, C. & Huson, D.H. Fast and sensitive protein alignment using DIAMOND. Nat. Methods 12, 59–60 (2015).

50. Galperin, M.Y., Makarova, K.S., Wolf, Y.I. & Koonin, E.V. Expanded microbial genome coverage and improved protein family annotation in the COG database. Nucleic Acids Res. 43, D261–269 (2015).

51. Overbeek, R. et al. The SEED and the Rapid Annotation of microbial genomes using Subsystems Technology (RAST). Nucleic Acids Res. 42, D206–214 (2014).

52. Letunic, I. & Bork, P. 20 years of the SMART protein domain annotation resource. Nucleic Acids Res. 46, D493–d496 (2018).

53. Couvin, D. et al. CRISPRCasFinder, an update of CRISRFinder, includes a portable version, enhanced performance and integrates search for Cas proteins. Nucleic Acids Res. 46, W246–w251 (2018).

54. Zhang, F. et al. CRISPRminer is a knowledge base for exploring CRISPR-Cas systems in microbe and phage interactions. Commun. Biol. 1, 180 (2018).

55. Emms, D.M. & Kelly, S. OrthoFinder: solving fundamental biases in whole genome comparisons dramatically improves orthogroup inference accuracy. Genome Biol. 16, 157 (2015).

56. Csuros, M. Count: evolutionary analysis of phylogenetic profiles with parsimony and likelihood. Bioinformatics 26, 1910–1912 (2010).

57. Katoh, K. & Standley, D.M. MAFFT multiple sequence alignment software version 7: improvements in performance and usability. Mol. Biol. Evol. 30, 772–780 (2013).

58. Capella-Gutierrez, S., Silla-Martinez, J.M. & Gabaldon, T. trimAl: a tool for automated alignment trimming in large-scale phylogenetic analyses. Bioinformatics. 25, 1972–1973. (2009).

59. Nguyen, L.T., Schmidt, H.A., von Haeseler, A. & Minh, B.Q. IQ-TREE: a fast and effective stochastic algorithm for estimating maximum-likelihood phylogenies. Mol. Biol. Evol. 32, 268–274 (2015).

60. Vaidya, G., Lohman, D.J. & Meier, R. SequenceMatrix: concatenation software for the fast assembly of multi-gene datasets with character set and codon information. Cladistics 27, 171–180 (2011).

