## Supplementary material for "Reductive evolution and unique infection and feeding mode in the CPR predatory bacterium *Vampirococcus lugosii*"

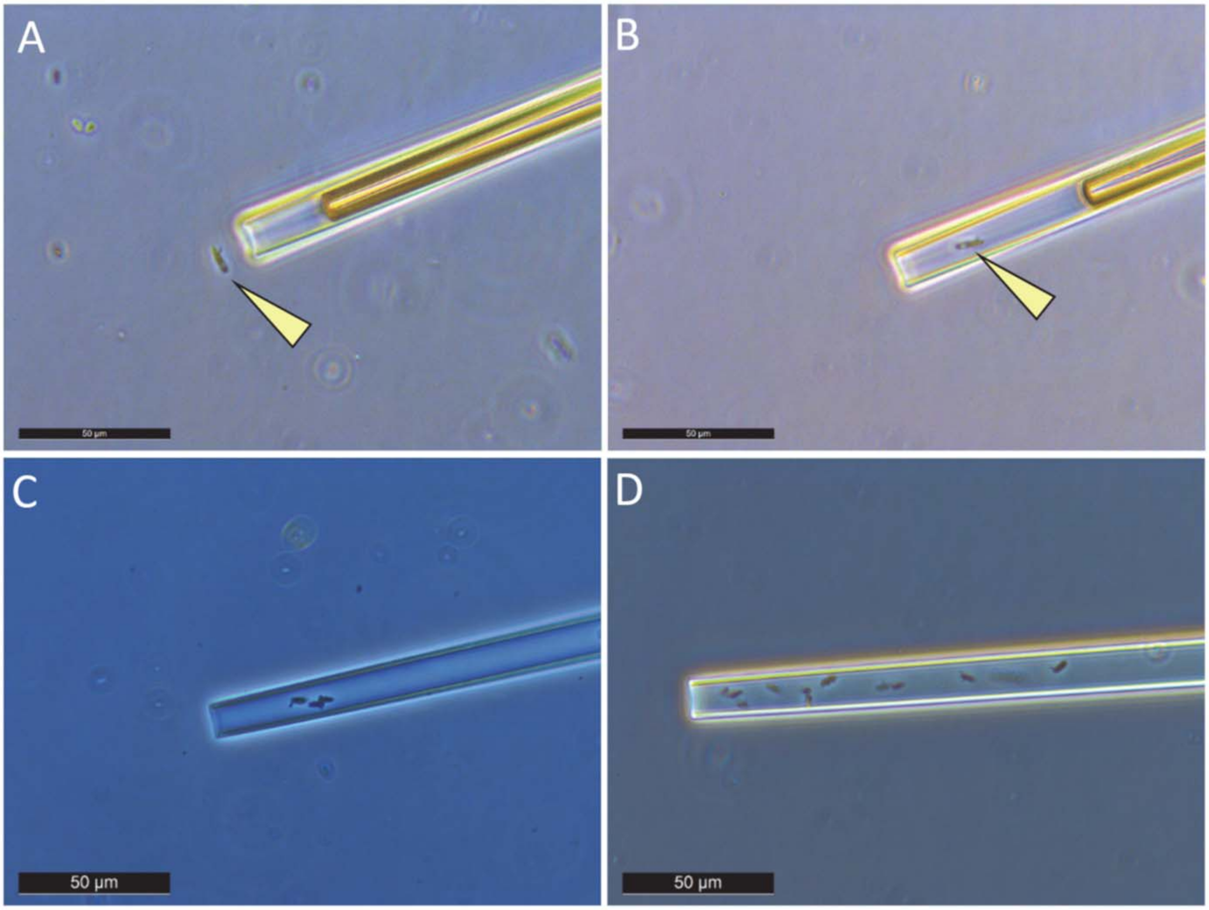


**Extended Data Fig. 1 |** Phase contrast microscopy images of micromanipulation of cells infected by *Vampirococcus* using a glass capillary. Yellow arrows in panels A and B indicate the same infected cells outside and inside the capillary. Panels C and D show capillaries containing several infected cells.

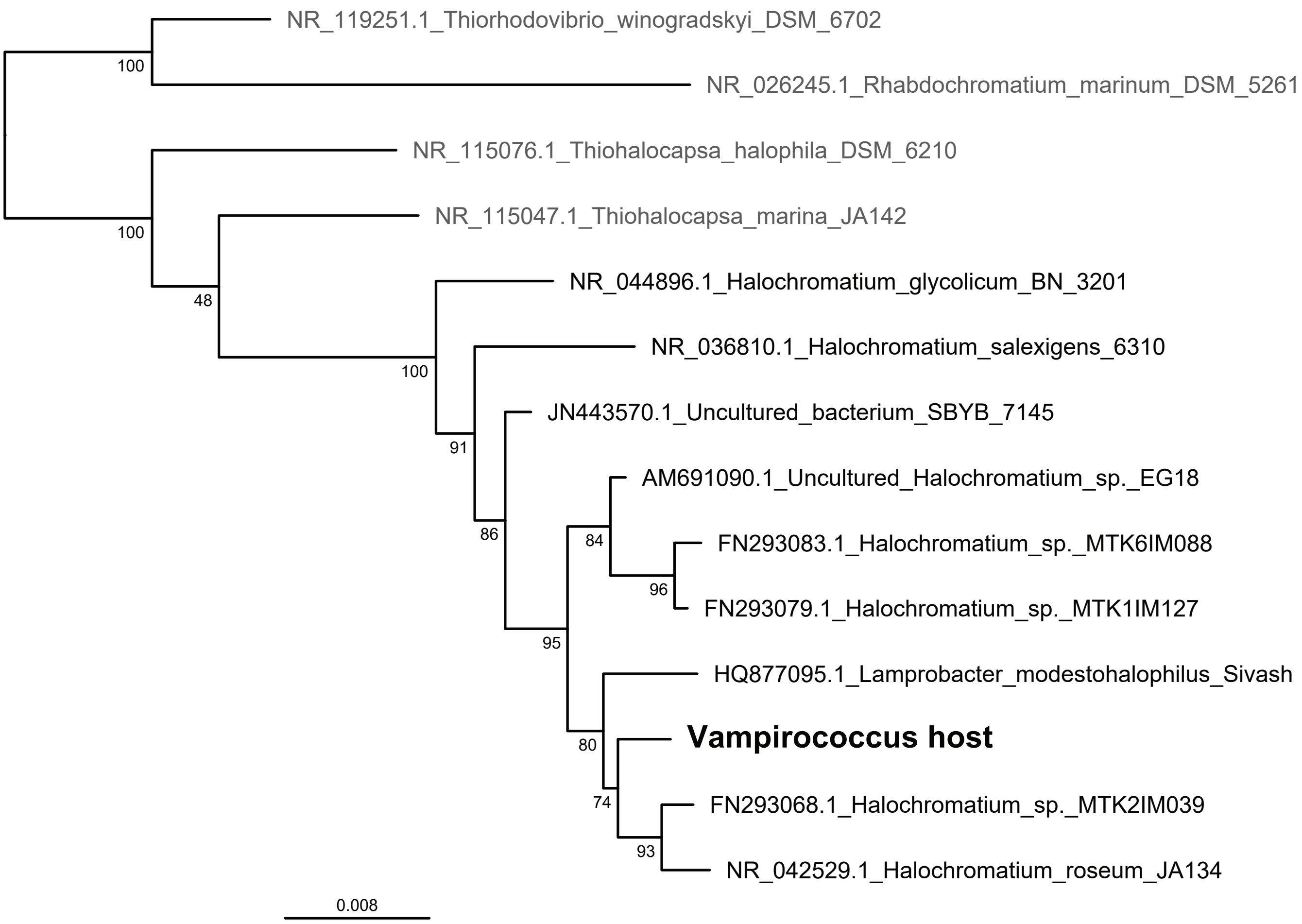


**Extended Data Fig. 2 |** 16S rRNA gene maximum likelihood phylogenetic tree of the *Vampirococcus* host. The tree is based on 1425 conserved aligned positions. Numbers at nodes are bootstrap support values (100 replicates).

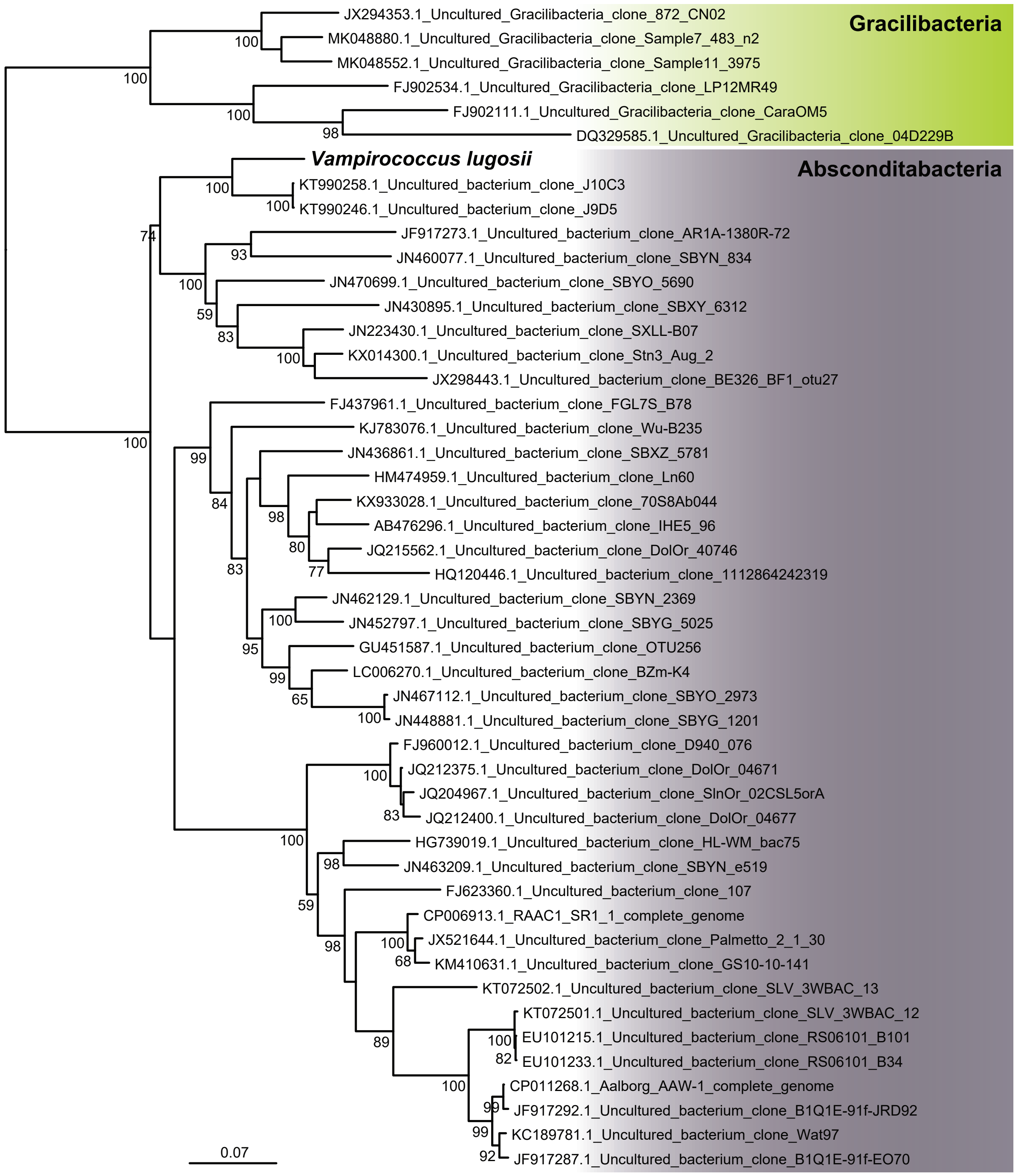


**Extended Data Fig. 3 |** 16S rRNA gene maximum likelihood phylogenetic tree of *Vampirococcus*. The tree is based on 1282 conserved aligned positions. Numbers at nodes are bootstrap support values (100 replicates). The tree is rooted on several Gracilibacteria sequences.

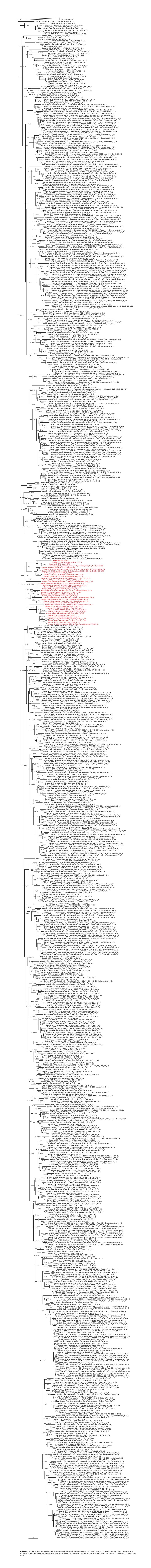

**Extended Data Fig. 4 |** Maximum likelihood phylogenetic tree of CPR bacteria showing the position of *Vampirococcus*. The tree is based on the concatenation of 16 ribosomal proteins and rooted on other bacteria. Numbers at nodes are bootstrap support values (100 replicates). The group containing *Vampirococcus* is indicated in red.

| Protein | Function |
| --- | --- |
| VAMP_2_16 | ATPase AAA |
| VAMP_2_102 | uracil-DNA glycosylase |
| VAMP_2_126 | amino acid kinase family protein |
| VAMP_2_142 | acid phosphatase family membrane protein YuiD |
| VAMP_2_172 | 5'(3')-deoxyribonucleotidase, containing HAD-like domain |
| VAMP_2_376 | coproporphyrinogen III oxidase |
| VAMP_2_652 | spermidine synthase |
| VAMP_2_667 | peptidase M50B family, putative |
| VAMP_2_731 | hypothetical protein |
| VAMP_5_14 | Asp-tRNAAsn/Glu-tRNAGln amidotransferase |
| VAMP_5_86 | arginyl-tRNA synthetase |
| VAMP_5_322 | sulfate permease |
| VAMP_5_352 | hypothetical protein |
| VAMP_6_101 | UDP-N-acetylenolpyruvoylglucosamine reductase |
| VAMP_7_1 | group 1 glycosyl transferase |
| VAMP_7_38 | glycosyltransferase |
| VAMP_7_63 | sulfotransferase |
| VAMP_7_102 | ATPase, AAA+ superfamily |
| VAMP_8_21 | putative bacterial transferase |
| VAMP_8_278 | zinc transporter ZupT |
| VAMP_11_1 | type III restriction endonuclease |
| VAMP_11_32 | death-on-curing protein |
| VAMP_11_87 | membrane protein |
| VAMP_11_209 | hypothetical protein |
| VAMP_12_204 | hypothetical protein |
| VAMP_12_314 | inosine 5'-monophosphate dehydrogenase |
| VAMP_12_407 | membrane protein |
| VAMP_12_450 | restriction endonuclease |
| VAMP_13_135 | adenosylhomocysteinase |
| VAMP_13_214 | lysophospholipase |
| VAMP_13_272 | membrane protein |
| VAMP_13_309 | ABC transporter |
| VAMP_13_360 | superoxide dismutase |
| VAMP_16_4 | hypothetical protein |
| VAMP_74-20_301 | metal-dependent hydrolase |
| VAMP_74-20_334 | type I site-specific deoxyribonuclease |
| VAMP_74-20_345 | helix-turn-helix domain-containing protein |
| VAMP_74-20_352 | hypothetical protein |
| VAMP_21_15 | CRISPR-associated protein Csm3 |
| VAMP_21_88 | CRISPR/Cas system-associated protein |
| VAMP_21_151 | CRISPR-associated protein Cas1 |
| VAMP_21_166 | CRISPR/Cas system-associated endoribonuclease |
| VAMP_22_222 | HD superfamily hydrolase (HD-GYP... |
| VAMP_24_237 | restriction endonuclease S subunit |
| VAMP_25_1 | HD superfamily hydrolase |
| VAMP_25_65 | putative diaminopimelate decarboxylase |
| VAMP_27_9 | ATPase AAA, partial |
| VAMP_27_16 | K+ transport system, NAD-binding |
| VAMP_27_59 | hypothetical protein |
| VAMP_27_173 | CDP-diglyceride synthetase |
| VAMP_31-433_4 | putative transposase |
| VAMP_34_196 | adenine specific DNA methylase |
| VAMP_34_248 | putative DEAD/DEAH box helicase |
| VAMP_34_317 | transposase |
| VAMP_36_45 | predicted ATPase |
| VAMP_36_80 | steroid 5-alpha reductase family |
| VAMP_36_96 | CAAX protease, self-immunity related |

| | |
|---|---|
| VAMP_40_134 | surface antigen |
| VAMP_40_177 | hAD phosphatase family IIIA |
| VAMP_41_97 | C4-dicarboxylate ABC transporter |
| VAMP_44_14 | thymidylate synthase |
| VAMP_44_34 | ribosome-associated GTPase EngA |
| VAMP_44_51 | helicase |
| VAMP_44_164 | transposase, IS4 family |
| VAMP_46_8 | ATPase AAA |
| VAMP_46_21 | restriction endonuclease |
| VAMP_46_157 | nitrous oxidase accessory protein |
| VAMP_48_7 | ATPase AAA |
| VAMP_48_145 | transposase |
| VAMP_48_161 | Kazal-type serine protease inhibitor |
| VAMP_68-56_291 | integrase |
| VAMP_59_23 | hypothetical protein |
| VAMP_62_140 | DNA methylase N-4 |
| VAMP_63_143 | hypothetical protein |
| VAMP_64_89 | aspartate racemase |
| VAMP_64_127 | integrase |
| VAMP_67_138 | DNA binding protein, containing HHH motif |
| VAMP_68-56_2 | hypothetical protein |
| VAMP_70_1 | transposase |
| VAMP_74-20_92 | hypothetical protein |
| VAMP_84_6 | transposase, partial |
| VAMP_84_29 | cell wall assembly/cell proliferation coordinating protein, KNR4 |
| VAMP_84_167 | transposase |
| VAMP_85_12 | ATPase |
| VAMP_85_59 | methyl-accepting chemotaxis protein (MCP) |
| VAMP_98_42 | ATP-dependent endonuclease |
| VAMP_98_79 | modification methylase HgaI-1 |
| VAMP_98_91 | type I restriction endonuclease subunit M |
| VAMP_98_96 | DNA-binding beta-propeller fold protein YncE |
| VAMP_102_119 | pectin lyase, putative |
| VAMP_106-108_174 | metallo-beta-lactamase superfamily protein |
| VAMP_115_23 | sialyltransferase, partial |
| VAMP_119_19 | DNA-binding protein |
| VAMP_119_36 | glycosyltransferase, GT2 family |
| VAMP_119_100 | DNA binding protein, containing KilA-N domain |
| VAMP_125_114 | hypothetical protein |
| VAMP_134_102 | NAD(P)-dependent dehydrogenase |
| VAMP_134_122 | AAA+ ATPase superfamily |
| VAMP_141_15 | membrane protein |
| VAMP_144_15 | nucleoside-triphosphatase |
| VAMP_152_39 | DNA methyltransferase |
| VAMP_163_8 | hypothetical protein |
| VAMP_163_14 | hypothetical protein |
| VAMP_163_27 | hypothetical protein |
| VAMP_163_38 | transcription factor E (TFE) |
| VAMP_163_45 | tRNA methyltransferase |
| VAMP_169_4 | 5-methylcytosine-specific restriction related enzyme |
| VAMP_197_9 | AAA+ ATPase superfamily |
| VAMP_237_13 | restriction endonuclease subunit S |
| VAMP_247_44 | protease, CAAX family |
| VAMP_247_64 | SPFH domain-containing protein |
| VAMP_283_8 | ATPase AAA |
| VAMP_315_21 | hypothetical protein |
| VAMP_325_1 | outer membrane protein OmpA |
| VAMP_328_89 | membrane protein |

| | |
|---|---|
| VAMP_368_47 | protein of unknown function, DUF285 |
| VAMP_379_21 | cold-shock protein |
| VAMP_397_11 | ribonuclease H |
| VAMP_405_17 | ParB-like and HNH nuclease domains |
| VAMP_31-433_257 | hypothetical protein |
| VAMP_472_45 | hypothetical protein |
| VAMP_493_19 | membrane protein |
| VAMP_525_55 | 5-methylcytosine-specific restriction endonuclease McrBC |
| VAMP_18_255 | ATPase AAA |
| VAMP_18_221 | chemotaxis protein |
| VAMP_18_122 | hypothetical protein |

**Extended Data Fig. 5 |** *Vampirococcus* genes acquired by horizontal gene transfer.

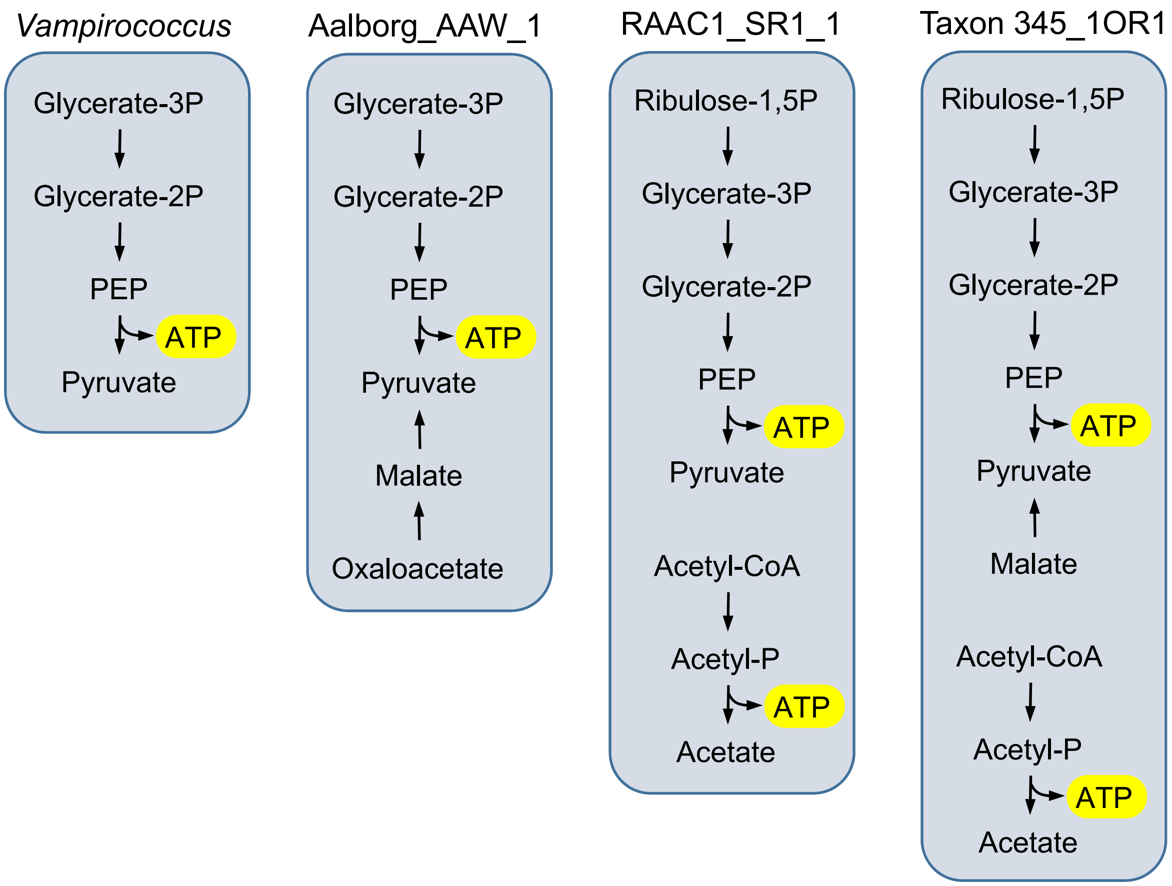


**Extended Data Fig. 6 |** ATP production pathways in *Vampirococcus* and three representative Absconditabacteria.

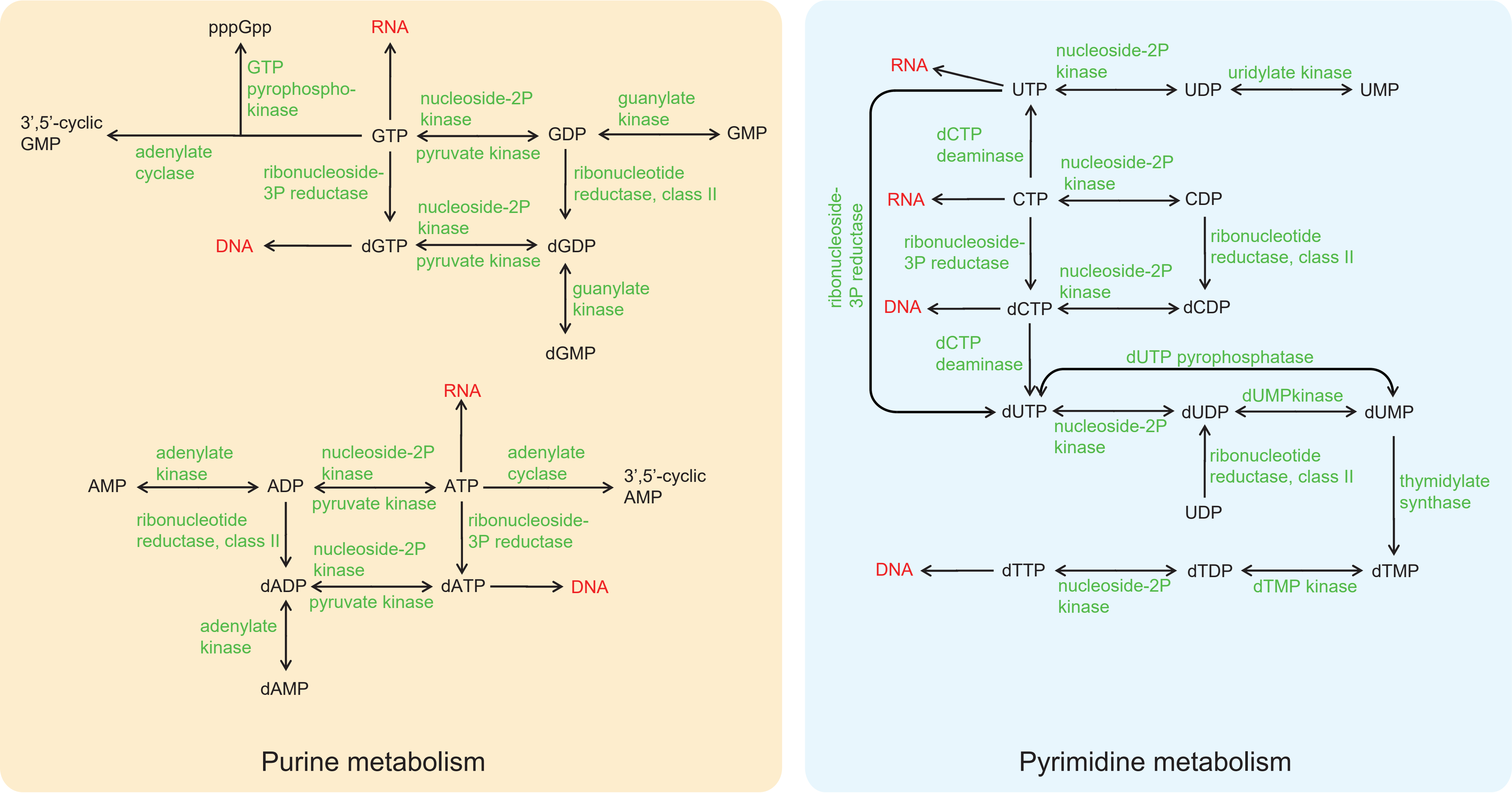


**Extended Data Fig. 7 |** Main pathways involved in purine and pyrimidine metabolism in *Vampirococcus*. Substrates are indicated in black, enzymes in green, and DNA and RNA in red.

| Protein | Function | Size |
|---|---|---|
| VAMP_16_20 | hypothetical protein | 4163 |
| VAMP_134_102 | NAD(P)-dependent dehydrogenase | 2400 |
| VAMP_6_203 | alpha-2 macroglobulin | 2368 |
| VAMP_166_2 | hypothetical protein | 2284 |
| VAMP_19_245 | alpha-2 macroglobulin | 1895 |
| VAMP_328_89 | membrane protein | 1883 |
| VAMP_95_65 | YfaS, alpha-2 macroglobulin family | 1871 |
| VAMP_54_69 | hypothetical protein | 1723 |
| VAMP_17-88-45_126 | PKD repeat-containing protein | 1396 |
| VAMP_98_96 | DNA-binding beta propeller protein YncE | 1392 |

**Extended Data Fig. 8 |** Ten largest proteins encoded in the *Vampirococcus* genome.

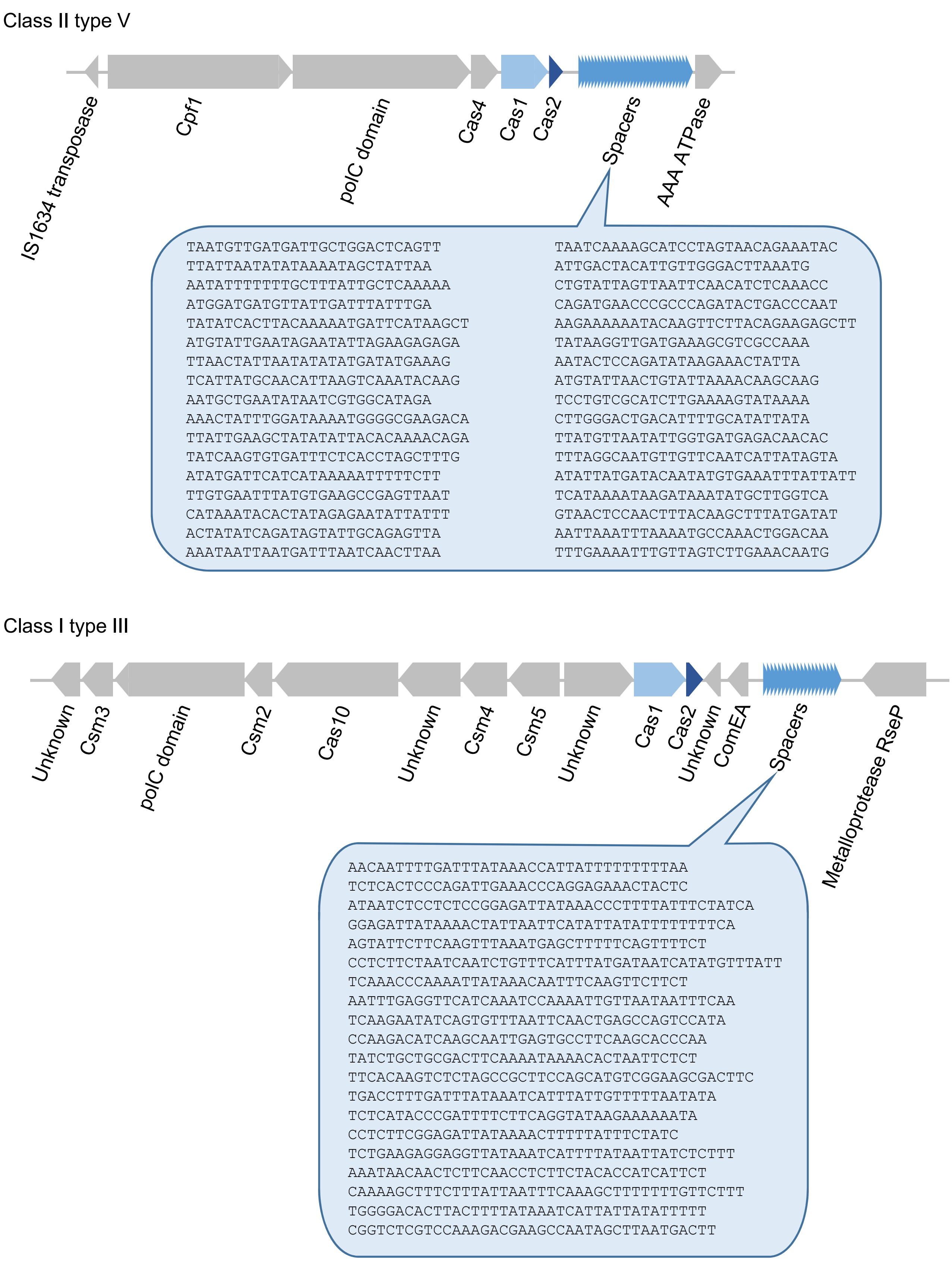


**Extended Data Fig. 9 | The two CRISPR-Cas loci of *Vampirococcus* and their corresponding spacer sequences.**

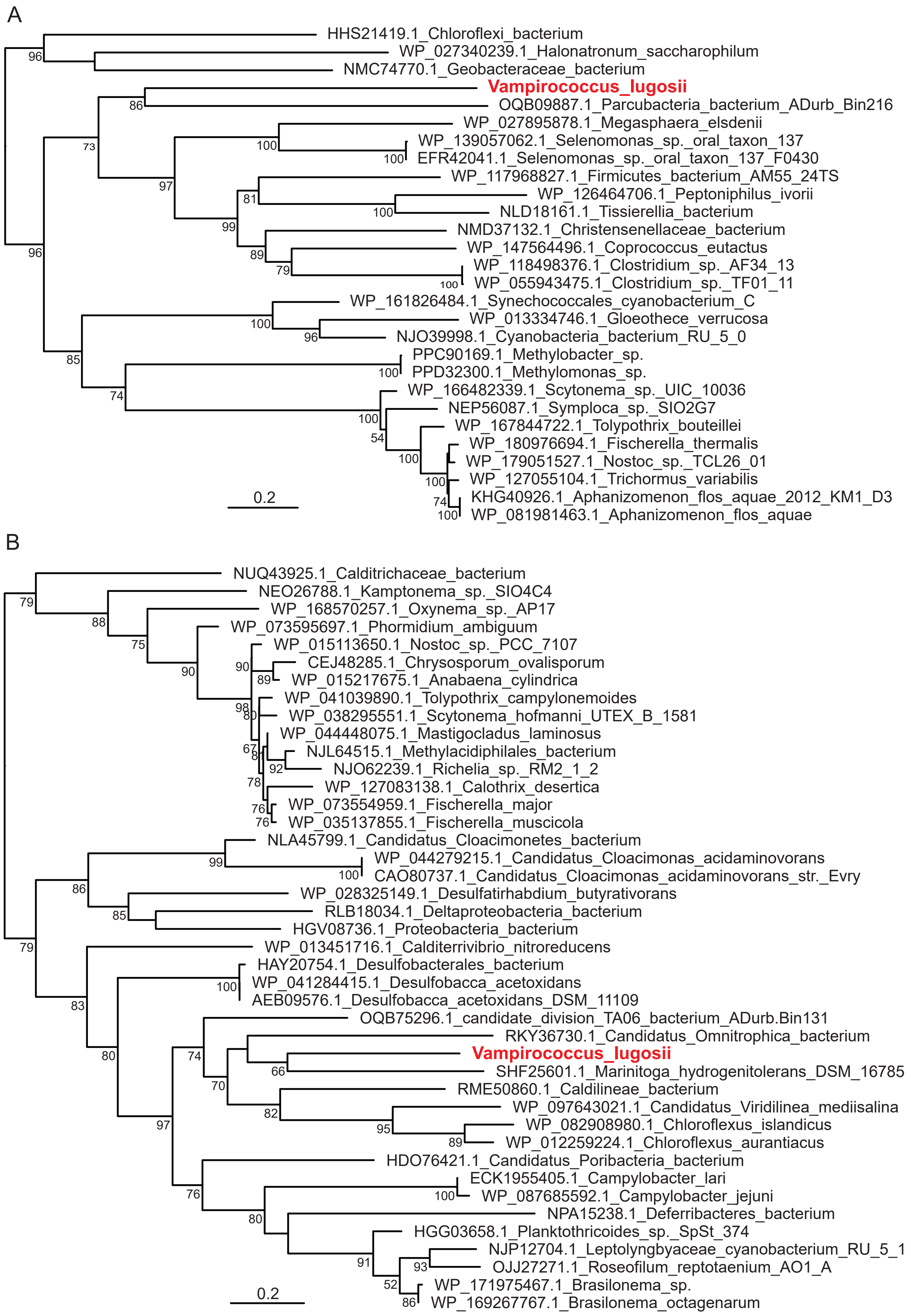


**Extended Data Fig. 10 |** Maximum likelihood phylogenetic trees of *Vampirococcus* type I class III CRISPR-Cas enzymes Cas1 ‘A) and Cas2 (B). The trees are based on 349 and 118 conserved aligned positions, respectively. Numbers at nodes are bootstrap support values (100 replicates, only values >50% are shown).

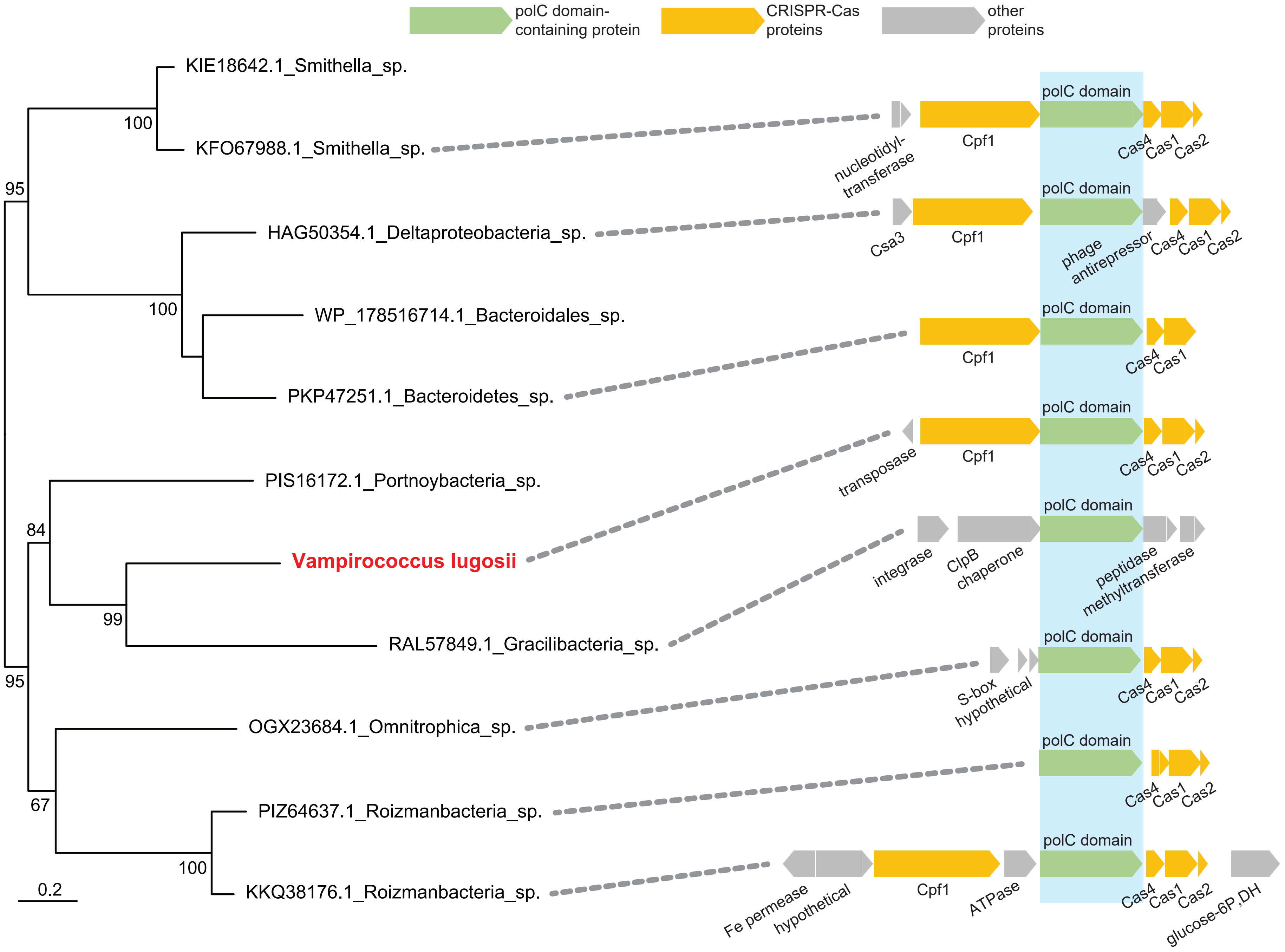


**Extended Data Fig. 11 |** Maximum likelihood phylogeny and genomic context of the *Vampirococcus* polC domain-containing protein associated to the class II type V CRISPR-Cas system. The genomic context is shown for the species for which genome sequence data are available. The polC domain-containing protein is shown in green and typical CRISPR-Cas proteins are shown in orange.

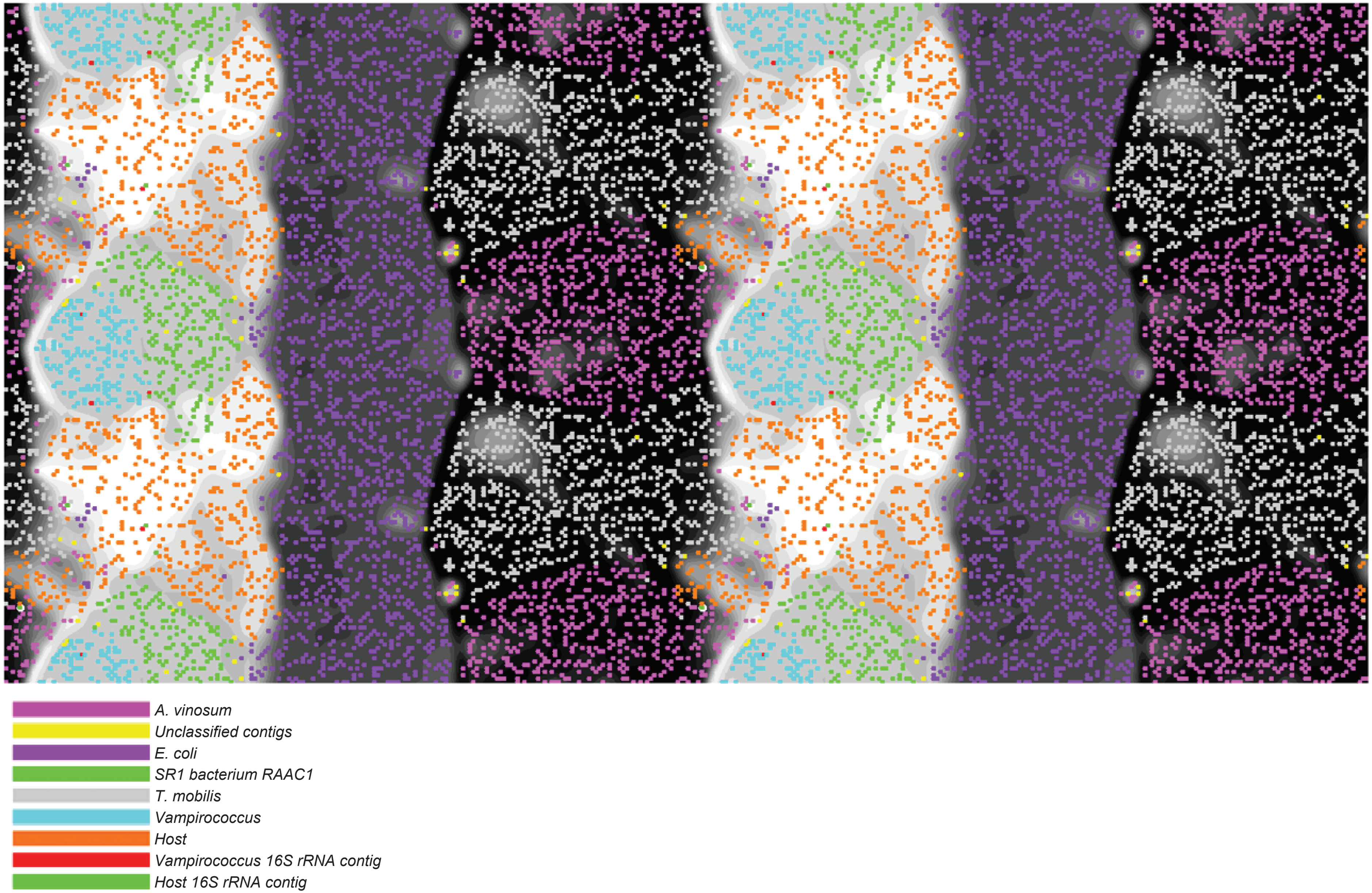


**Extended Data Fig. 12 |** Visualization of the ESOM map of the assembled DNA contigs of *Vampirococcus*, its host, and several reference genomes.
